# ABC for high-dimensional modular models via MCMC samples

**DOI:** 10.1101/2025.07.02.662793

**Authors:** Zhixiao Zhu, Maria Christodoulou, David Steinsaltz

## Abstract

Many complex systems are modelled using *modular models*, where individual sub-models are estimated separately and then combined. While this simplifies inference, it fails to account for interactions between components. A natural solution is to estimate all components jointly, but this is often impractical due to intractable likelihoods. Approximate Bayesian Computation (ABC) provides a likelihood-free alternative, but its standard implementations are computationally inefficient, particularly when applied to high-dimensional modular models, or when sub-models involve costly machine learning methods, like Gaussian Process (GP) models. The ABC-Population Monte Carlo (ABC-PMC) framework improves on vanilla ABC by using sequential Monte Carlo sampling with adaptive tolerances and proposal kernels, yielding much higher acceptance rates and more efficient exploration of parameter space. Existing ABC-PMC algorithms are not, however, especially efficient in the high-dimensional parameter setting typical of modular models.

We introduce a novel modification of the ABC-PMC method that leverages model modularity. Our approach refines the prior distribution and perturbation kernel by using precomputed Markov Chain Monte Carlo (MCMC) samples from individual sub-models, making parameter updates more efficient. Additionally, we employ an adaptive summary statistic weighting strategy that dynamically adjusts the contribution of different statistics, reducing the influence of less informative statistics. These modifications greatly reduce overall computational cost. In our case studies, the runtime for 10,000 simulation attempts drops from over 20 days to under 1 minute, following a one-off preprocessing step that consists of standard MCMC sampling for each sub-model (typically 3-10 hours, depending on model complexity).

We apply our method to an ecological case study using an Integral Projection Model (IPM) for *Cryptantha flava*, where survival, growth, and reproduction processes are modelled using GP models. The results of the simulated and the real case studies demonstrate greatly improved computational efficiency while preserving inference quality. While the case study focuses on ecology, the method is applicable to a broad range of modular models where capturing interactions among sub-models is essential.

## 1 Introduction

Giant systems have become increasingly common across scientific research (Caswell (2000); Siciliano et al. (2008); Ellner et al. (2016); Figueiredo et al. (2018); Sato (2023)). To study these complex systems, researchers often use *modular models*, breaking down the system into smaller sub-models, each representing a specific process within the larger system. This approach typically starts with independent fitting of each sub-model, then combines them to gain a holistic understanding of the system as a whole. This “fit individually, then combine” approach is prevalent across various fields:

- In robotic systems, accurate manipulation requires precise calibration of various subsystems. It involves initially using simpler models to calibrate basic components, followed by more complex models to refine the calibration and capture intricate interactions between the robot’s mechanical and sensory systems (Sato (2023)).
- In energy system simulations, models are often fitted for multiple years using historical data. Each year’s data is used to fit individual linear regression models. These models are then combined into a multi-year model that accounts for annual variations, such as changes in weather conditions and other systemic factors over time (Figueiredo et al. (2018)).
- In biology, to understand population dynamics over time, models combine various vital rate sub-models, such as survival, growth, and reproduction, each fitted independently using field data. The sub-models are then combined to project the population structure over time, accounting for the interactions between different life stages (Caswell (2000); Ellner et al. (2016)).

While widely used, the “fit individually, then combine” approach has notable limitations. The main challenge lies in its assumption that accurately modeling each component will yield a reliable representation of the entire system. When fitted separately, sub-models do not account for interactions that naturally arise. Such interactions — arising from dependencies between processes or responses to shared conditions — can lead to system-level behaviours that isolated model fitting fails to consider. Models lacking such joint structure may therefore not fully capture the complexity of those giant systems.

Joint modeling of all modules within a single, fully combined framework might potentially capture these dependencies. However, constructing such a comprehensive “giant model” is generally infeasible. Complex systems with (potentially) many interacting components often lack well-defined likelihood functions, making it challenging to specify a unified framework representing the entire system. Without clear likelihoods and manageable computational demands, creating a fully combined model becomes impractical.

Approximate Bayesian Computation (ABC) provides a powerful alternative for statistical inference when likelihood functions are intractable or expensive to compute. These methods have widespread applications across various fields, including ecology, genetics, and epidemiology (Tavaré et al. (1997); Beaumont et al. (2002); Marjoram et al. (2003)). Instead of evaluating likelihoods directly, ABC methods approximate the posterior distribution by simulating data from the model and only retaining parameter values that are able to ‘reproduce’ observations. This makes ABC particularly well suited for modular models, where interactions among sub-models affect system behaviour in ways that are difficult to characterize explicitly. By selecting parameter values that enable the model to replicate observed data, ABC can help reveal interactions that influence system dynamics, even when they are not directly specified in the model structure. In these cases, inference is not only about estimating parameters but also about characterizing how different components jointly contribute observed outcomes.

However, ABC methods face challenges when applied to high-dimensional models (Sisson & Fan (2018)), as in the case of modular models. Traditional ABC approaches, including rejection sampling, ABC-MCMC, ABC-SMC and ABC-PMC (Tavaré et al. (1997); Marjoram et al. (2003); Cappé et al. (2004); Beaumont et al. (2009); Toni et al. (2009); Sisson et al. (2009)) struggle with high rejection rates and inefficient exploration of parameter space. The “curse of dimensionality” becomes an obstacle, as identifying perturbation kernels appropriate for all sub-models remains challenging. Inappropriate kernels can dramatically increase computational costs due to high rejection rates, especially as modern machine learning techniques, like Gaussian process (GP) models (Williams & Rasmussen (2006)), are increasingly adopted to model components of complex systems (Nabavi et al. (2023); Chen et al. (2024)). These models often contain intricate dependencies and latent structures across multiple dimensions, making it more challenging to design an effective global perturbation kernel.

In response to these challenges, we introduce a novel enhancement to the ABC-PMC algorithm, tailored specifically for modular models. Our method uses pre-calculated Markov Chain Monte Carlo (MCMC) samples from individual modules to initialise the prior distribution and design module-specific perturbation kernels. This guides exploration toward regions with high marginal posterior density, reducing the number of unlikely proposals and, thereby, lowering computational cost.

To further improve sampling efficiency, we incorporate an adaptive summary statistic weighting strategy proposed by Prangle (2017). This approach combines selected summary statistics into a single ‘universal’ distance metric, preventing less informative statistics from over-affecting the entire system. Overall, our approach considerably decreases the computational demands commonly encountered in high-dimensional settings, making ABC methods more viable for modular models.

This paper is structured as follows: in Section 2, we first provide an introduction to the foundational concepts of ABC methods; then, we detail our version of ABC-PMC and the specific modifications we have implemented. In section 3, we demonstrate its utility through a well-established population model, the Integral Projection Model (IPM), applied to the shrub *Cryptantha flava*, where flexible GP models are used for sub-models of vital rates. The results of the simulated and real case studies demonstrate greatly improved computational efficiency while maintaining inference reliability within a modular model framework. Section 4 discusses the method’s applicability and future research directions.

## 2 ABC algorithms

ABC has been progressively refined to enhance computational efficiency and sampling performance (e.g. Tavaré et al. (1997); Marjoram et al. (2003); Wegmann et al. (2009); Beaumont et al. (2009); Toni et al. (2009); Sisson et al. (2009); Lenormand et al. (2013); Andrieu et al. (2018)). This section presents key approaches, beginning with ABC rejection sampling as a baseline, followed by ABC importance sampling, and finally introducing ABC population Monte Carlo (ABC-PMC) as an iterative extension.

Consider Bayesian inference for a parameter vector ***θ*** *∈* Θ under a model with the likelihood *f* (*y*_obs_|***θ***) with the observed dataset *y*_obs_ *∈ 𝒟*. Given a prior distribution *π*(***θ***), the posterior distribution *π*(***θ***|*y*_obs_) *∝ f* (*y*_obs_|***θ***)*π*(***θ***). We further assume that evaluating the likelihood *f* (*y*_obs_|***θ***) is computationally costly or numerically impossible, but generating simulations from the model is relatively straightforward.

### 2.1 ABC rejection sampler

A basic ABC approach (Algorithm 1), introduced by Tavaré et al. (1997), extends standard rejection sampling. It proceeds by sampling parameter candidates ***θ***\* from the prior, simulating datasets *y*\* under each sampled parameter, and accepting those that exactly reproduce the observed dataset, *y*\* = *y*_obs_.

#### Algorithm 1: ABC Rejection sampling method

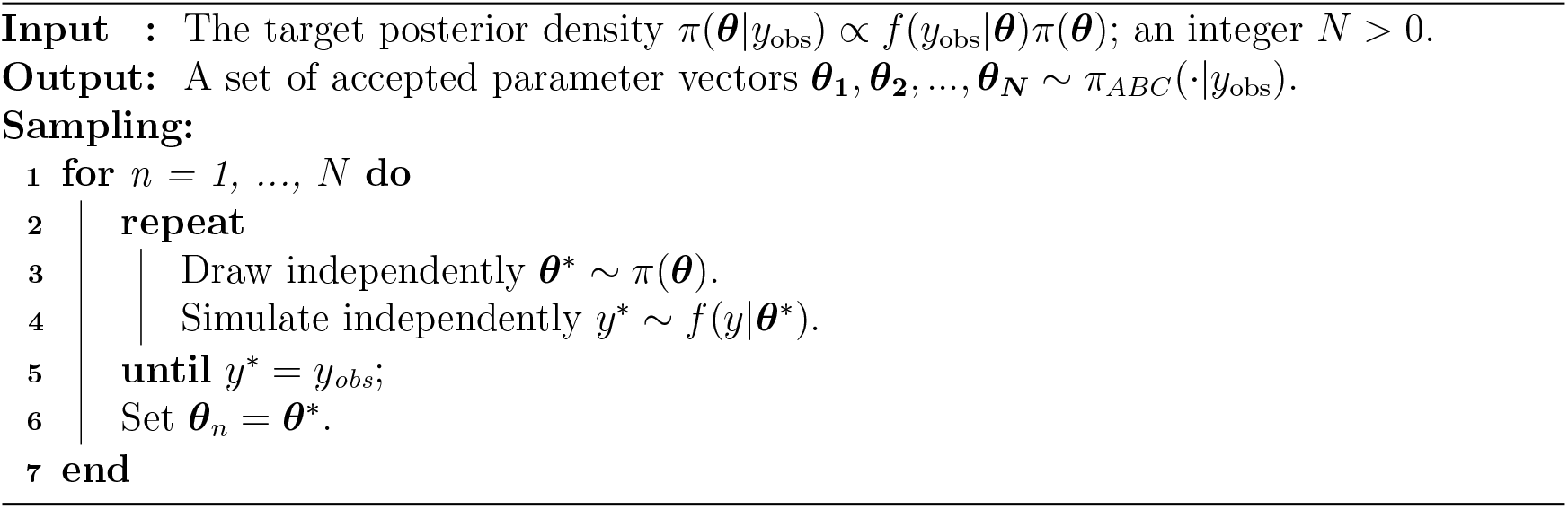

This method theoretically ensures the accepted parameters from the exact posterior, i.e. *π*_*ABC*_(***θ***|*y*_obs_) = *π*(***θ***|*y*_obs_), where *π*_*ABC*_(***θ***|*y*_obs_) denotes the approximate posterior obtained through ABC sampling. However, requiring exact dataset replication is impractical for most applications, particularly when data is continuous, as the acceptance probability approaches zero (Sisson et al. (2018)). To improve feasibility, a typical implementation of ABC replaces the full dataset *y*_obs_ with a low-dimensional vector of summary statistics 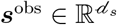. Instead of requiring an exact match, the algorithm accepts ***θ***\* if the simulated summary ***s***\* is sufficiently close to ***s***^obs^, measured by distance functions ***d***(***s***^obs^, ***s***\*), like the Euclidean distance (Fu & Li (1997); Wall (2000); Tishkoff et al. (2001)). A tolerance threshold ***h*** *>* 0 controls the acceptance region, with acceptance occurring when ***d***(***s***^obs^, ***s***\*) *<* ***h***. When applying these modifications to Algorithm 1, line 5 is replaced by *d*_*i*_(^*i*^*s*^*\*, i*^*s*^obs^) *< h*_*i*_ for *i* in 1, …, *d*_*s*_, where *d*_*i*_ and *h*_*i*_ denote the distance metric and acceptance criterion for the *i*th summary statistic respectively. Thus, a parameter is accepted only if all *d*_*s*_ summary statistics satisfy their respective thresholds.

As a result, ABC samplers rely on the assumption that *π*(***θ***|*y*_obs_) can be reasonably approximated by *π*(***θ***|***s***^obs^). Given this assumption, ABC objective is, instead, to approximate the distribution *π*(***θ***|***s***^obs^), where lim_***h****→*0_ *π*_*ABC*_(***θ***|***s***^obs^) = lim_***h****→*0_ *π*_*ABC*_(***θ***|***d***(***s***\*, ***s***^obs^) *<* ***h***) = *π*(***θ***|***s***^obs^). A natural way to achieve this is by considering the joint distribution of ***θ*** and ***s***\*, and marginalizing over ***s***\*, i.e. *π*(***θ***|***s***^obs^) = *π*(***θ, s***\*|***s***^obs^)*d****s***\* (Sisson & Fan (2018)). Especially, in rejection-based ABC, the joint distribution *π*(***θ, s***\*|***s***^obs^) is approximated by

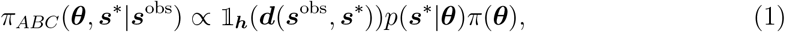

where 𝟙_***h***_(*·*) is an indicator function that determines whether the distance ***d***(***s***^obs^, ***s***\*) is within the acceptance tolerance ***h***. For simplicity, we use 𝟙_***h***_(*·*) here, but it can be replaced by more general smoothing kernels, see Peters et al. (2012) for example.

### 2.2 ABC importance sampler

The efficiency of ABC rejection sampling is greatly affected by the choice of prior *π*(***θ***). In real-world applications, the posterior is often quite different from the prior. In such cases, proposed values from *π*(***θ***) can frequently fall in low-probability regions of the posterior (Marin et al. (2012)), leading to high rejection rates and inefficient inference. A natural extension is to introduce a proposal distribution *q*(***θ***) that can better focus on high-probability posterior regions, rather than relying solely on the prior. To ensure coverage of the posterior, we require *q*(***θ***) *>* 0 whenever *π*(***θ***|***s***^obs^) *>* 0. This modification leads to the ABC importance sampler. Accordingly, the approximation in Equation 1 becomes

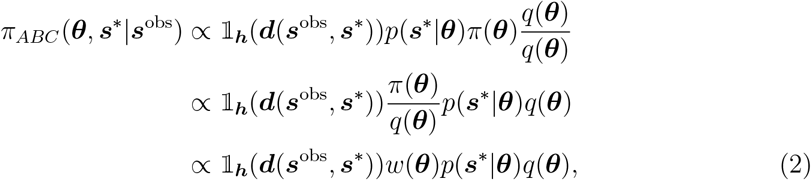

where the importance weight is defined as *w*(***θ***) = *π*(***θ***)*/q*(***θ***). Thus, ABC importance sampling retains the rejection-based structure but incorporates a correction scheme through importance weighting.

### 2.3 ABC-PMC

While importance sampling improves efficiency, it relies on a fixed proposal *q*(***θ***). Choosing an appropriate *q*(***θ***) can be challenging (Marin et al. (2012)), particularly when the posterior distribution is highly concentrated or multimodal. A poorly chosen proposal may either be too broad, leading to excessive sampling in low-probability regions, or too narrow, failing to sufficiently explore the posterior. In such cases, relying on a single fixed, pre-specified proposal can be highly limited, especially since the posterior shape is generally unknown.

One might wonder: given a set of accepted parameter values, can we leverage these samples to refine our proposal distribution dynamically, ensuring that future proposals better align with high-probability regions of the posterior? This idea forms the basis of ABC-PMC (Beaumont et al. (2009); Toni et al. (2009); Sisson et al. (2009)).

ABC-PMC (Algorithm 2) builds on importance sampling by iteratively refining the proposal distribution. At each iteration, it generates a population of *N* accepted particles, and then adjusts the proposal distribution *q*_*c*_(***θ***) by favouring well-performing particles from previous iterations while discarding lower-weighted ones (Beaumont et al. (2009); Toni et al. (2009)). This process progressively refines the approximation to the target posterior, via a sequence of slowly evolving intermediate distributions.

When implementing ABC-PMC only once, *C* = 0, it is nothing more than a standard ABC importance sampler. When *C >* 0, ABC-PMC is in essence an iterative version of the ABC importance sampler (Cappé et al. (2004); Lintusaari et al. (2017)). That is (to align with Equation 2), for each iteration, *c* = 0, …, *C*, ABC-PMC draw samples from the target

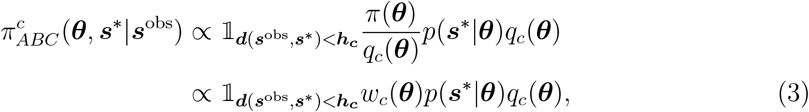

where bandwidth ***h***_***c***_ is a decreasing sequence (***h***_**0**_ *> · · · >* ***h***_***C***_ ⩾ 0), so that each successive intermediate target can be less diffuse and gradually converge to the final target distribution, 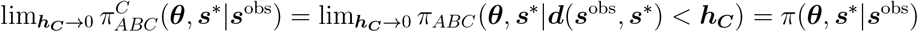 (Toni et al. (2009); Sisson & Fan (2018)).

#### Algorithm 2: Conventional ABC-PMC

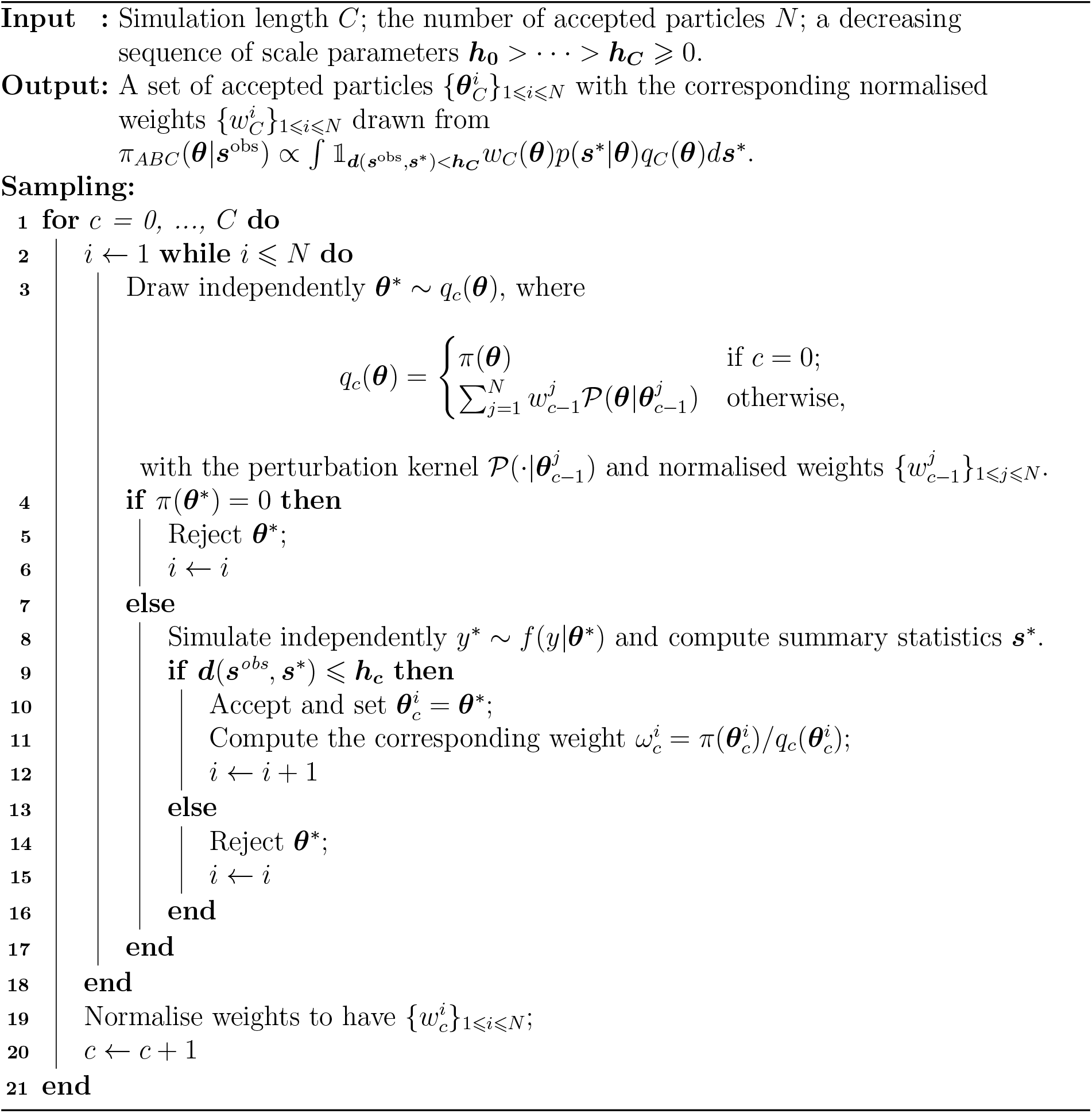

### 2.4 Modifications to Conventional ABC-PMC

ABC-PMC improves ABC importance samplers by iteratively refining the proposal distribution, but its efficiency fundamentally depends on the choice of *perturbation kernels P* (Line 3 in Algorithm 2). Conventional ABC-PMC methods typically rely on, for example, component-wise uniform (Barnes et al. (2011)), component-wise normal (Beaumont et al. (2009)), or multivariate normal perturbation kernels (Filippi et al. (2013)). In modular models, interactions between and within sub-models introduce complex dependencies, making it difficult for standard perturbation kernels to propose effective updates: component-wise kernels ignore these dependencies; while multivariate normal kernels capture only linear correlations (Khazeiynasab & Qi (2021)).

To address these inefficiencies, we introduce a novel sampling framework that replaces conventional perturbation kernels with a Gibbs-like, MCMC-informed proposal mechanism. Instead of perturbing all parameters simultaneously, we randomly select one sub-model at each iteration and update all of its parameters using pre-computed MCMC posterior samples. This targeted update strategy preserves dependencies within each sub-model while accelerating the proposal process.

Specifically, in our case studies, the MCMC pre-computation takes approximately 3–10 hours per sub-model (depending on model complexity). These runs are independent across sub-models and can be parallelised. Once MCMC samples are pre-computed and stored, the computational time for generating 10,000 sampling *proposals* (not *acceptances*) is reduced by a factor of around 10^4^ — from around 20 days to under 1 minute in our example. The developed version of the ABC-PMC method, detailed in Algorithm 3 and Table 1, operates as follows:

**Table 1:** The description of notation used in Algorithm 3.

| Notation | Description |
| --- | --- |
| $\mathcal{M}$ | The target modular model. |
| $\top$ | The transpose of a matrix. |
| $C$ | The total number of simulation steps for ABC-PMC (apart from the initialisation). |
| $c$ | The current simulation step; $c = 0$ for initialisation. |
| $N_c$ | The number of accepted particles required for the simulation step $c$. |
| $\alpha_c$ | The quantile employed to decide the acceptance region for the next simulation $c + 1$. |
| $V$ | The total number of sub-models in the target modular model. |
| $v$ | A realisation of a discrete uniformly distributed random variable indicating the current sub-model. |
| $M$ | The number of MCMC samples for each of the $V$ sub-models. So, there are $M^V$ different combinations in total. |
| $m_v$ | Given the current sub-model $v$, $m_v$ is a realisation of a discrete uniformly distributed random variable representing $m_v$-th MCMC sample for $v$th sub-model. |
| $\theta$ | A particle: a vector including parameters for all of the sub-models. |
| $\theta_c^i$ | The $i$th particle at $c$th simulation step. |
| $\tilde{\theta}$ | An intermediate particle. $\tilde{\theta}^\top = (\tilde{\theta}_1^\top, \dots, \tilde{\theta}_V^\top)$, where $\tilde{\theta}_v$ for $v = 1, \dots, V$ is a vector only includes $v$th sub-model's parameters. |
| $\tilde{\theta}^*$ | A perturbed intermediate particle. |
| $\pi(\theta)$ | The prior for $\theta$, which is the joint distribution for $m_1, \dots, m_V$. |
| $\omega_c^i$ | Unnormalised weight for $\theta_c^i$ |
| $w_c^i$ | Normalised weight for $\theta_c^i$ |
| $y^*$ | A simulated dataset. |
| $y_{\text{obs}}$ | The observed dataset. |
| $d_s$ | Dimension of $\mathbf{S}_c^i$ and $\tilde{\mathbf{S}}^*$. |
| $\mathbf{S}_c^i$ | $\mathbf{S}_c^i = ({}^1S_c^i, {}^2S_c^i, {}^3S_c^i, {}^4S_c^i, \dots, {}^{d_s}S_c^i)$, where $({}^jS_c^i)$ is $j$th summary statistics for $i$th particle at step $c$. |
| $\tilde{\mathbf{S}}^*$ | A vector of summary statistics produced by $\tilde{\theta}^*$, $\tilde{\mathbf{S}}^* = ({}^1\tilde{S}^*, \dots, {}^{d_s}\tilde{S}^*)$. |
| $\sigma_c$ | A vector of MADs for all of the selected summary statistics, $\sigma_c = ({}^1\sigma_c, \dots, {}^{d_s}\sigma_c)$. |
| $d_c^i$ | $d_c^i = (\sum_{j=1}^{d_s} ({}^jS_c^i / {}^j\sigma_c)^2)^{1/2}$, which represent the ‘universal’ distance between $y_{\text{obs}}$ and a simulated dataset produced by $\theta_c^i$. |
| $\tilde{d}^*$ | $\tilde{d}^* = (\sum_{j=1}^{d_s} ({}^j\tilde{S}^* / {}^j\sigma_{c-1})^2)^{1/2}$. |
| $\delta_c$ | $\alpha_c$th quantile of $\{d_c^i\}_{1 \leq i \leq N_c}$; the quantile-based acceptance threshold at step $c$. |
| $\mathbb{1}_\tau$ | An indicator function, $\mathbb{1}_\tau = 1$ if the statement $\tau$ is true; $\mathbb{1}_\tau = 0$ otherwise. |
| $a \rightarrow b$ | This is a statement about whether it is possible to transit from state $a$ to state $b$ by the perturbation kernel at line 15 in Algorithm 3. |

#### 1 Initialisation

Start with particles, drawn from an *informative* prior based on pre-calculated MCMC samples, simulating datasets and calculating initial distances using summary statistics. Set initial weights for all particles equally.

#### 2. Iterative Process

- For each step *c* = 1 to *C*, resample particles based on their weights from step *c* − 1 and perturb them by randomly altering one sub-model’s parameter using MCMC samples.
- Simulate a new dataset from each perturbed particle and recalculate the distance to the observed data.
- Accept particles if their recalculated distance is below a *dynamically* computed quantile-based threshold. Update and normalise weights of accepted particles.

#### 3. Output

Continue until the final iteration, producing a weighted set of particles that approximate the posterior distribution.

Compared to the conventional ABC-PMC method (Algorithm 2), we highlight three key modifications in our method (Algorithm 3):

1. **We employ a bespoke prior distribution** *π*(*·*) **and perturbation kernel** *𝒫*. After obtaining MCMC samples for each sub-model individually — to gain insights into the posterior distributions of their own parameters, we then
  - *π*(*·*): re-use the MCMC samples for each individual sub-model to build up an *informative* prior for the entire big model. Specifically, after obtaining, for example, *M* different MCMC samples for each of *V* individual sub-models, the resulting informative prior has a support (sample space) containing *M*^*V*^ possible points.
  - 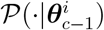: Given a particle 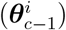 accepted in the previous iteration, a new movement is proposed by randomly selecting one of the *V* sub-models and replacing all corresponding parameters with a randomly chosen sample from the *M* MCMC samples (Line 15 in Algorithm 3).
2. Illustrated by the toy example in Appendix A, through an adaptive weighting strategy proposed by Prangle (2017), **the selected summary statistics are combined into a single ‘universal’ distance** to measure the difference between the observed and simulated dataset. In summary, the ‘universal’ distance between the observed and a simulated dataset at the *c*-th iteration is defined by

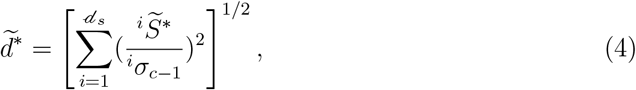

where ^*i*^*σ*_*c*−1_ is the estimated median absolute deviation (MAD) (based on the (*c* − 1)-th generation of particles) for the *i*-th summary statistic 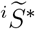. Notice that, for simplicity, we refer to 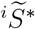 as a ‘summary statistic’, though it actually represents the distance between summary statistics from the simulated and observed datasets. We will maintain this notation throughout the report.
3. Lastly, conventional ABC-PMC (Algorithm 2) leaves the choice of bandwidth *{****h***_***c***_*}*_*c*=0, …, *C*_ up to the users themselves, but it could be difficult to specify or tune it well: large values lead to poor posterior approximations, while small values reduce acceptance rates and increase computational cost. A widely used adaptive approach, introduced by Drovandi & Pettitt (2011), sets *{****h***_***c***_*}* to be the *α* quantile of the accepted distances at the previous iteration. However, using a constant quantile *α* in Algorithm 3 brings us a potential risk that the determined acceptance regions might not belong to subsets of a nested sequence converging to the point 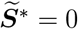 (Schälte et al. (2021)).

We consider various strategies for defining acceptance regions, ultimately settling on **a decreasing sequence of quantiles** *{α*_*i*_*}*_0⩽*i*⩽*C*−1_. To balance computational time, an additional modification is made by **gradually reducing total number of accepted particles required**, *{N*_*i*_*}*_0⩽*i*⩽*C*_. A proper balance between these two factors enable us to propagate only the most promising particles, providing stability for the future samples in Algorithm 3. Table 2 provides practical examples of *{α*_*i*_*}* and *{N*_*i*_*}* used in the case study.

**Table 2:** Results of applying ABC-PMC to IPM in a simulated dataset. **Notations** : *δ*_**c**−**1**_: the quantile-based acceptance threshold used at step *c*, which was determined based on *α*_**c**−**1**_ quantile at step *c* _−_ 1 (so denoted by *δ*_*c*−1_). *δ*_−1_ and *α*_−1_ are for the initialisation. 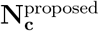: the proposed number of accepted particles required for the simulation step *c*. **N**_**c**_: the true number of accepted particles for the simulation step *c*. 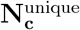: the number of unique particles among *N*_*c*_ acceptances. **n**(***𝒳***_**c**_): the total number of possible states of ***θ*** at simulation step *c*. Apart from **n**(***𝒳***_**0**_), the others were estimated through *n*(***𝒳***_*c*_)^*lb*^. 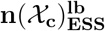 : a lower bound for *n*(***𝒳***_*c*_) but estimated based of ESS. 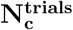: total number of trials to achieve the acceptance. 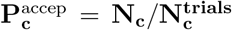: the acceptance rate for step *c*. 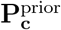: the acceptance rate for a single-iterated ABC-PMC, where the samples were drawn directly from the priors made up by MCMC samples. 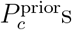 were computed based on 10,000,290 particles drawn from *π*(***θ***). To ensure comparability, 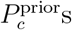 must be computed under the exactly same setting that produced 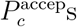. That is, for each *c* in *{*0, 1, …, *C}*, the 10,000,290 particles were normalised with MADs ***σ***_*c*−1_ first, the threshold *δ*_*c*−1_ then employed to compute 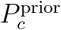. It took approximately 530.5 minutes to complete this single-iterated algorithm with 10,000,290 particles. **t**_**c**_: the total amount of time, measured in minutes, used to complete the *c*th iteration.

| | $c = 0$ | $c = 1$ | $c = 2$ | $c = 3$ | $c = 4$ |
| --- | --- | --- | --- | --- | --- |
| $\alpha_{c-1}$ | 1 | 0.50 | 0.50 | 0.45 | 0.40 |
| $\delta_{c-1}$ | ∞ | 10.82 | 9.70 | 9.00 | 8.41 |
| $N_c^{\text{proposed}}$ | 1,000,000 | 500,000 | 300,000 | 150,000 | 100,000 |
| $N_c$ | 1,000,060 | 501,875 | 300,176 | 150,650 | 100,435 |
| $N_c^{\text{unique}}$ | 1,000,060 | 501,869 | 300,173 | 150,650 | 100,435 |
| $n(\mathcal{X}_c)$ | 5000<sup>5</sup> | $> 2.5 \times 10^{10}$ | $> 1.1 \times 10^{10}$ | $> 6.5 \times 10^9$ | $> 3.3 \times 10^9$ |
| $n(\mathcal{X}_c)_{ESS}^{lb}$ | 5000<sup>5</sup> | $> 2.5 \times 10^{10}$ | $> 1.1 \times 10^{10}$ | $> 6.5 \times 10^9$ | $> 3.2 \times 10^9$ |
| $N_c^{\text{trials}}$ | 1,000,060 | 1,000,000 | 1,070,000 | 1,040,000 | 1,500,000 |
| $P_c^{\text{accep}}$ | 1 | 0.5019 | 0.2805 | 0.1449 | 0.0670 |
| $P_c^{\text{prior}}$ | 1 | 0.4999 | 0.2502 | 0.1101 | 0.0422 |
| $t_c$ | 53.15 | 75.17 | 77.78 | 74.55 | 107.97 |

Table 2: Results of applying ABC-PMC to IPM in a simulated dataset.
| | $c = 5$ | $c = 6$ | $c = 7$ | $c = 8$ | $c = 9$ |
| --- | --- | --- | --- | --- | --- |
| $\alpha_{c-1}$ | 0.35 | 0.30 | 0.20 | 0.15 | 0.10 |
| $\delta_{c-1}$ | 7.92 | 7.38 | 6.78 | 6.19 | 5.56 |
| $N_c^{\text{proposed}}$ | 60,000 | 30,000 | 15,000 | 7,000 | 1,000 |
| $N_c$ | 60,040 | 30,036 | 15,026 | 7,003 | 1,000 |
| $N_c^{\text{unique}}$ | 60,039 | 30,036 | 15,026 | 7,002 | 1,000 |
| $n(\mathcal{X}_c)$ | $> 2.1 \times 10^9$ | $> 1.2 \times 10^9$ | $> 6.4 \times 10^8$ | $> 3.2 \times 10^8$ | $> 1.5 \times 10^8$ |
| $n(\mathcal{X}_c)_{ESS}^{lb}$ | $> 2.0 \times 10^9$ | $> 1.2 \times 10^9$ | $> 6.0 \times 10^8$ | $> 2.9 \times 10^8$ | $> 1.3 \times 10^8$ |
| $N_c^{\text{trials}}$ | 2,190,000 | 3,120,000 | 6,370,000 | 16,740,000 | 19,980,000 |
| $P_c^{\text{accep}}$ | 0.0274 | 0.0096 | 0.0024 | $4.183 \times 10^{-4}$ | $5.005 \times 10^{-5}$ |
| $P_c^{\text{prior}}$ | 0.0139 | 0.0038 | 0.0007 | $8.600 \times 10^{-5}$ | $8.300 \times 10^{-6}$ |
| $t_c$ | 158.12 | 225.1 | 453.22 | 1182.57 | 1433.43 |

For more detailed explanations about these modifications please refer to Appendix 2.4. The appendix describes the motivations and thought process behind the modifications, providing insights for those looking to apply or extend our approach.

It is worth noting that, although both the sub-model MCMC inference and the ABC posterior approximation rely on the same underlying dataset, they draw on different levels of information. The sub-models are typically fitted using individual-level observations — like covariates, transitions, or outcomes associated with individual components — whereas the ABC step compares simulated and observed summary statistics defined at the system or population level. These summary statistics, like event rates or empirical distributions across groups, represent aggregate summaries of the data that do not retain individual-level structure. As a result, the information used in sub-model MCMC fitting is not directly reused in the ABC sampling. While this setup does not constitute a fully independent use of data, we consider the two stages to target distinct aspects of the dataset. In this sense, it differs from typical cases in which the same information is reused twice.

##### Algorithm 3: ABC-PMC (see Table 1 for the notation)

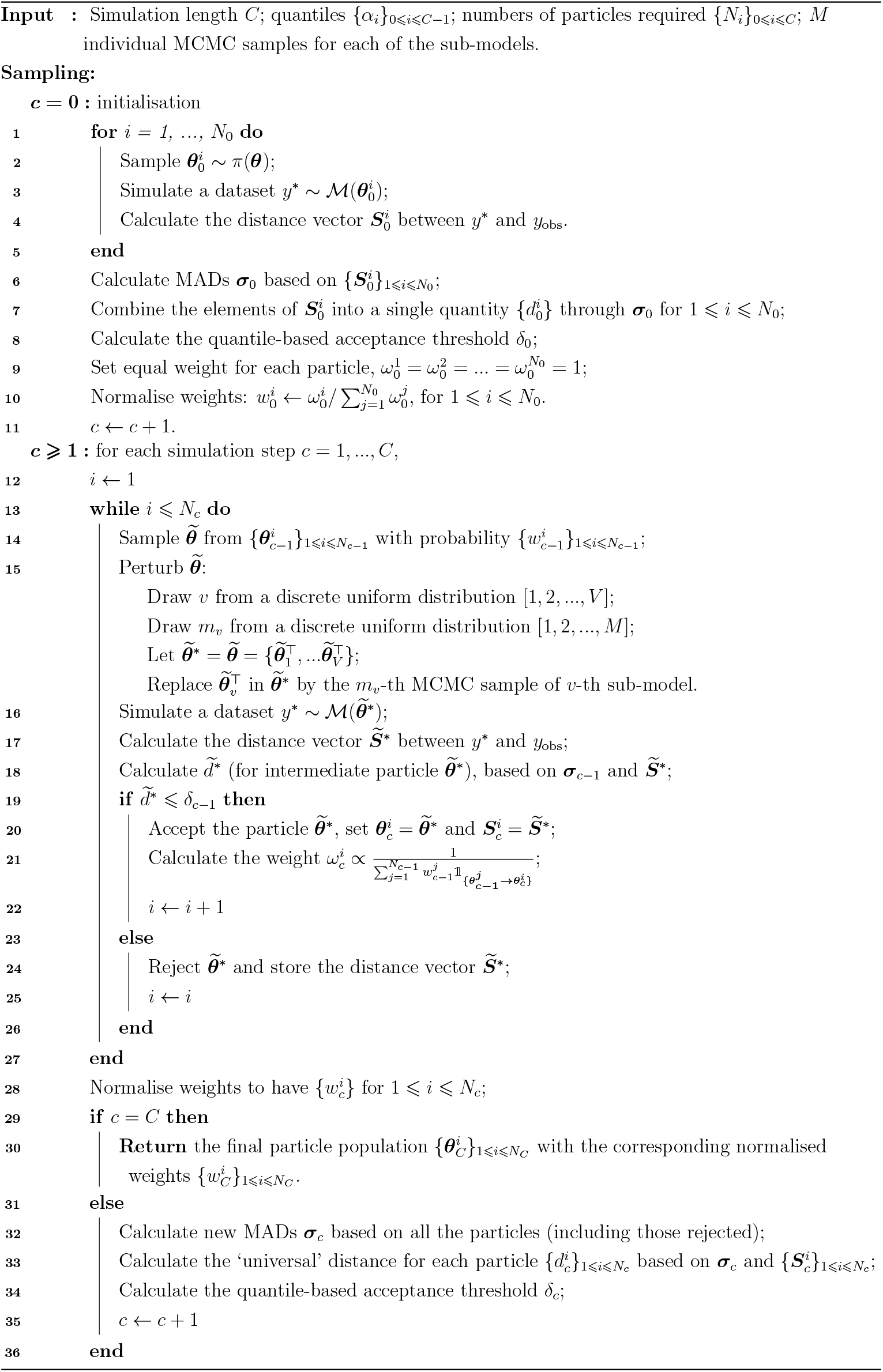

## 3 Case studies

In this section, we applied our modified ABC-PMC sampler to Integral Projection Models (IPMs) for *Cryptantha flava*. To assess its effectiveness, we first conducted a case study using simulated datasets with known ground truth. With each iteration, the estimated population growth rate increasingly aligned with the true rate, validating the method’s reliability. Building on this, we extended the ABC-PMC sampler to the real datasets of *C. flava*.

Initially, we intended to compare our results with traditional ABC-PMC methods, but this was computationally infeasible. In our experiments, each simulation attempt took approximately 3 minutes to run on a Apple M1 Max processor. The simulation had to proceed sequentially, as later stages of the simulation depend on earlier outcomes, for example, growth depends on whether an individual survives. At this rate, 10,000 simulation attempts would take at least 20 days — 10,000 acceptances are insufficient for one iteration in conventional ABC-PMC, let alone for just 10,000 attempts.

In contrast, our method requires a one-off preprocessing step: running MCMC for each sub-model independently. In our experiments, this took around 3–10 hours per sub-model depending on complexity, but the runs can be performed in parallel. After this step, the runtime for 10,000 sampling attempts drops to under 1 minute in our case studies.

Due to the impractical runtime of standard ABC-PMC, we were unable to conduct a benchmark comparison. Our evaluation thus focuses solely on the proposed method. The fundamental components of the model’s framework are outlined below, with full experiment details available in Zhu et al. (2025).

### 3.1 IPM overview and Experimental setup

An IPM is considered a modular model that combines various sub-models describing biological processes like survival, growth into a single framework to project population dynamics (Ellner et al. (2016)). This approach allows researchers to simultaneously consider multiple factors that influence population dynamics. IPMs assume that the individuals in the target population can be completely summarised by an attribute *z* (like the individual size) (Rees et al. (2006); Ellner & Rees (2006)). In IPMs, with current attribute *z*, a population at the next time step with attribute *z*′ can be described by

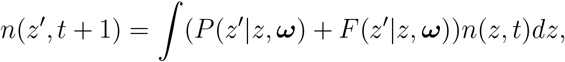

where ***ω*** represents environmental factors affecting the *survival-growth kernel, P* (*z*′|*z*, ***ω***), and the *reproduction kernel, F* (*z*′|*z*, ***ω***).

To reflect *C. flava*’s life cycle, *P* and *F* can be further separated into individual vital rates to align with the actual life cycle events. *F* (*z*′|*z*, ***ω***) starts with considering the probability of breeding, *p*_*f*_ (*z*|***ω***); conditional on flowering, a state *z* individual would produce an average of *n*_*f*_ (*z*|***ω***) flowering stalks; each stalk leads to a certain count of seeds establishing and growing to size *z*′ by the next census, *r*_*est*_*r*_*size*_(*z*′). That is, *F* (*z*′|*z*, ***ω***) = *p*_*f*_ (*z*|***ω***)*n*_*f*_ (*z*|***ω***)*r*_*est*|***ω***_*r*_*size*_(*z*′|***ω***). The survival-growth kernel, *P* (*z*′|*z*, ***ω***), considers the reproduction’s influence on growth dynamics. It assumes that an individual with current state *z* will survive to the next year with probability *s*(*z*|***ω***), but the resulting growth would depend on whether or not the individual reproduces: *G*_*f*_ (*z*′|*z*, ***ω***) or *G*_*nf*_ (*z*′|*z*, ***ω***). That is, *P* (*z*′|*z*, ***ω***) = *s*(*z*|***ω***)*p*_*f*_ (*z*|***ω***)*G*_*f*_ (*z*′|*z*, ***ω***) + *s*(*z*|***ω***)(1 − *p*_*f*_ (*z*|***ω***))*G*_*nf*_ (*z*′|*z*, ***ω***). The sequence of life cycle events in *F* (*z*′|*z*, ***ω***) and *P* (*z*′|*z*, ***ω***) align with the actual census process conducted in Salguero-Gomez et al. (2012).

These sub-models of vital rates are parametrised by GP models. Traditional IPMs fit sub-models independently before combining them. Our approach, however, incorporates additional population-level information during integration (as in Zhu et al. (2025)). Since the likelihood of these additional population-level information is unknown, we employ the ABC method.

For the simulation study, we first fit GP models under a frequentist framework using population data from 2008 to 2011. We then used these fitted models to run Individual-based model (IBM) simulations to project population dynamics over a decade, creating a simulated dataset. This resulting dataset served as surrogate ‘truths’ for subsequent analysis.

For applying the sampler to real-world datasets of *C. flava*, model training used data from 2003 to 2008, with the last four years reserved for testing.

### 3.2 Results

#### 3.2.1 The simulation study

Table 2 provides a detailed summary of algorithm efficiency for the developed ABC-PMC. As expected, with the modified sequence of quantiles *{α*_*i*_*}*_0⩽*i*⩽10_, the acceptance threshold *δ*_*i*_ gradually decreases over the entire simulation process. Since computations were executed in parallel, the actual number of accepted particles (*N*_*c*_) sometimes slightly exceeded the proposed value 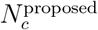. To maintain consistency with Algorithm 3, we use *N*_*c*_ to denote the true number of accepted particles.

#### Proposal Diversity

A key observation is that the number of unique accepted particles remains close to the total number of accepted particles 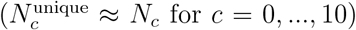. This may suggest that the algorithm does not frequently propose the same particles, avoiding redundant sampling.

Other observations from Table 2 further support this interpretation In our ABC-PMC framework, the proposal space at iteration *c* depends on the set of accepted particles from the previous iteration. Each accepted particle can generate up to 5*M* possible states for the next iteration. However, not all accepted particles contribute equally to exploration in the next iteration. Some particles tend to generate highly similar or even identical proposals, due to the employed perturbation kernel that preserve local structure. To describe this, we define 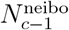 as the number of accepted particles that can transition to states that are already well-covered in the proposal space. These particles do not add diversity to the new proposals. To approximate the number of genuinely new proposal states, *n*(***𝒳***_*c*_), we estimate a lower bound for it as:

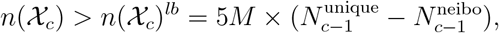

where 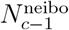 is estimated by the number of accepted particles within 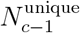 unique particles that share identical states in *V* −1 out of *V* sub-models with another accepted unique particle.

Additionally, since ABC-PMC operates as an iterative ABC importance sampling, another lower bound can be roughly estimated using the effective sample size (ESS):

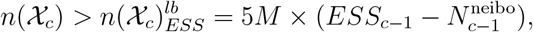

with 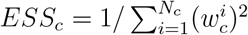.

From Table 2, we observe that the total number of trials required to obtain the desired number of acceptances is much smaller than the estimated size of the proposal space, i.e., 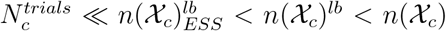 for *c* = 0, …, 10. Together with the previous observation that 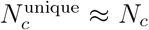 for *c* = 0, …, 10, there is not much space left for the algorithm to repeatedly propose the same particles (which lie in low posterior probability regions) multiple times and yield only a few acceptances among the repeated proposals.

#### Proposal Quality Assessment

If the distribution of the ‘universal’ distance remained unchanged throughout the ABC-PMC model fitting process, then running ABC-PMC with a sequence of decreasing acceptance thresholds *{δ*_*c*_*}*_0⩽*c*⩽*C*_ is nothing better than conducting a single iteration with the strictest threshold *δ*_*C*_. To evaluate the effectiveness of the proposed ABC-PMC, we examine its ability to improve proposal quality over iterations.

To check if there is an improvement, we compare the true acceptance rate 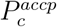 with 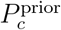, the acceptance rate of a single-iteration ABC-PMC using threshold *δ*_*c*−1_. To ensure comparability, 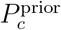 is computed under exactly the same setting used to generate 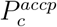 (see details in the caption of Table 2). In this case, if the proposals offer no advantage over the prior, then we expect 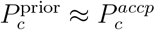 or even 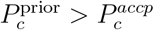.

The results from Table 2 and Figure 1 (Left) indicate that the proposal distribution, *q*_*c*_ evolved gradually to be much more efficient throughout the ABC-PMC. Initially, the ratio 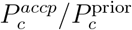 is close to 1, but it rapidly increases as the iterations proceed. By the end of the simulation, the true acceptance rate, 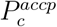, exceeds 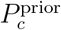 by a factor of more than 6.

**Figure 1:**
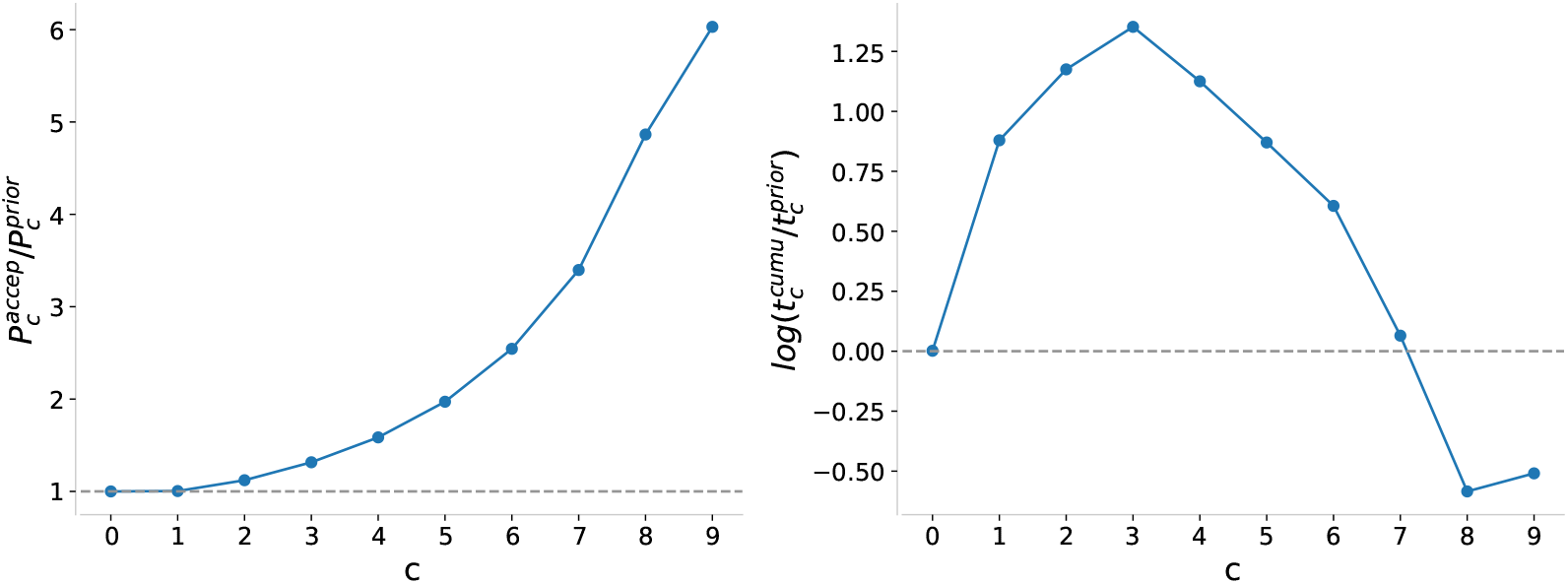
**Left:** The ratio of 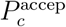 and 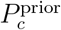 over simulation time *c*. The grey horizontal line cross the point representing 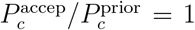. **Right:** The logarithmic ratio between the cumulative time spent on ABC-PMC, represented as 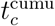, and the expected implementation time, estimated as 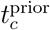. The 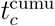 is obtained by adding up individual times for each single iteration *t*_*c*_ until the current iteration 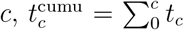. Instead of timing in real practices, 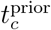was estimated using the formula 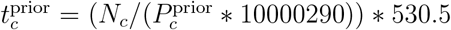 based on the 10,000,290 particles (see Table 2).

However, a higher acceptance rate in the proposal distribution does not necessarily imply that the ABC-PMC is more efficient. Before we get to the final round of proposals, we have incurred the computational cost of all the previous iterations that refined the distribution, so we need to sum up the time spent on the entire implementation.

This total cost is displayed on the right-hand side of Figure 1, plotting the log ratio between the cumulative runtime for ABC-PMC, 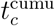, and the estimated runtime for a single iteration, 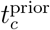, where the samples are drawn directly from the priors under the identical acceptance thresholds. Although the particles generated by ABC-PMC always exhibit higher acceptance probabilities, obtaining the same number of acceptances can initially require much more computational time. For example, at *c* = 3, ABC-PMC takes nearly four times longer than a single-iteration approach using MCMC priors (as 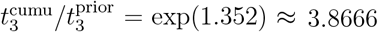). However, after *c* = 7, the proposed ABC-PMC becomes increasingly efficient. By the final iteration, the total runtime of ABC-PMC is only around 60% of the estimated runtime required for the single-iterated algorithm using just MCMC priors 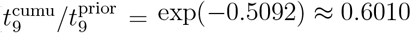 times).

#### Trends in Summary Statistics

Figure 2 illustrates the variations in the summary statistics throughout the ABC-PMC process. Looking at the orange lines, we see that the means (dashed) and medians (solid) of the 5 selected summary statistics gradually decrease as the acceptance threshold *δ*_*c*_ becomes more restrictive. This observation indicates the effectiveness of the adaptive weighting strategy, as the imposed barrier on the ‘universal distance’ can validly constrain each of the 5 summary statistics. Since the orange lines only reflect accepted particles, their decreasing pattern is expected, aligning with the setup of *{α*_*c*_*}*.

**Figure 2:**
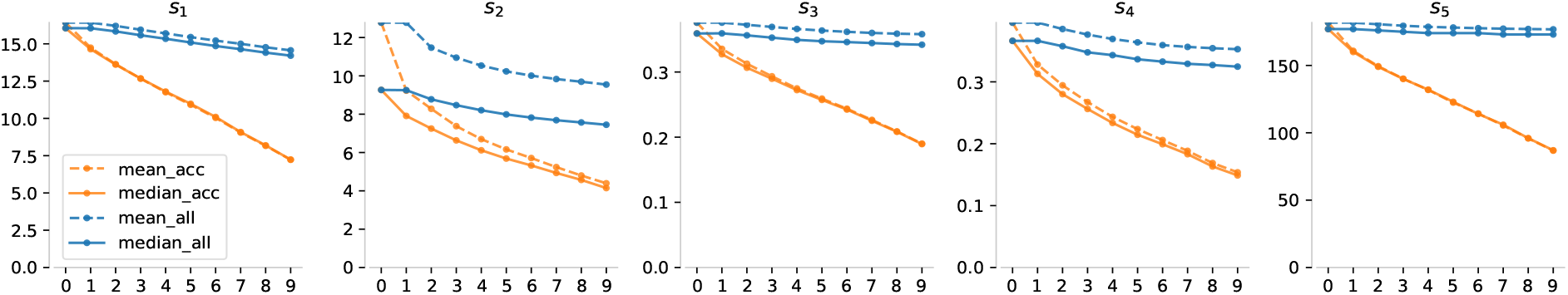
The mean (dashed line) and median (solid line) for the 5 selected summary statistics, *s*_1_ to *s*_5_, over the ABC-PMC simulation time *c* (*x* − axis). The orange lines describing the information of summary statistics for accepted particles only, while the blues stand for all particles including those rejected.

Conversely, the blue lines reveal a different perspective. They represent the summary statistics for all proposed particles (before any decisions to accept or reject are made). These particles can reflect the distance between the posterior *π*(***θ***|*y*_*obs*_) and the current proposal *q*_*c*_. If Figure 1 is not clear in demonstrating that the proposals *q*_*c*_ are continually enhanced throughout the algorithm, the descending trend of the blue lines in Figure 2 offers direct evidence of this improvement.

Given this understanding, it is not surprising to see in Figure 3 that, the predicted distribution of population growth rate (*λ*) was corrected step by step through a sequence of slowly varying intermediate distributions in each ABC-PMC iteration — even though we do not directly target *λ*. Note that, at step *c* = 10 of Figure 3, where we applied the most restrictive threshold, leading to an extremely high rejection rate and a limited yield of only 120 samples within a feasible time frame. Due to these constraints, the results from this step were excluded from our evaluations of predictive performance and algorithm efficiency. We present the findings about *c* = 10 here to illustrate the potential of the proposed method for achieving even better results.

**Figure 3:**
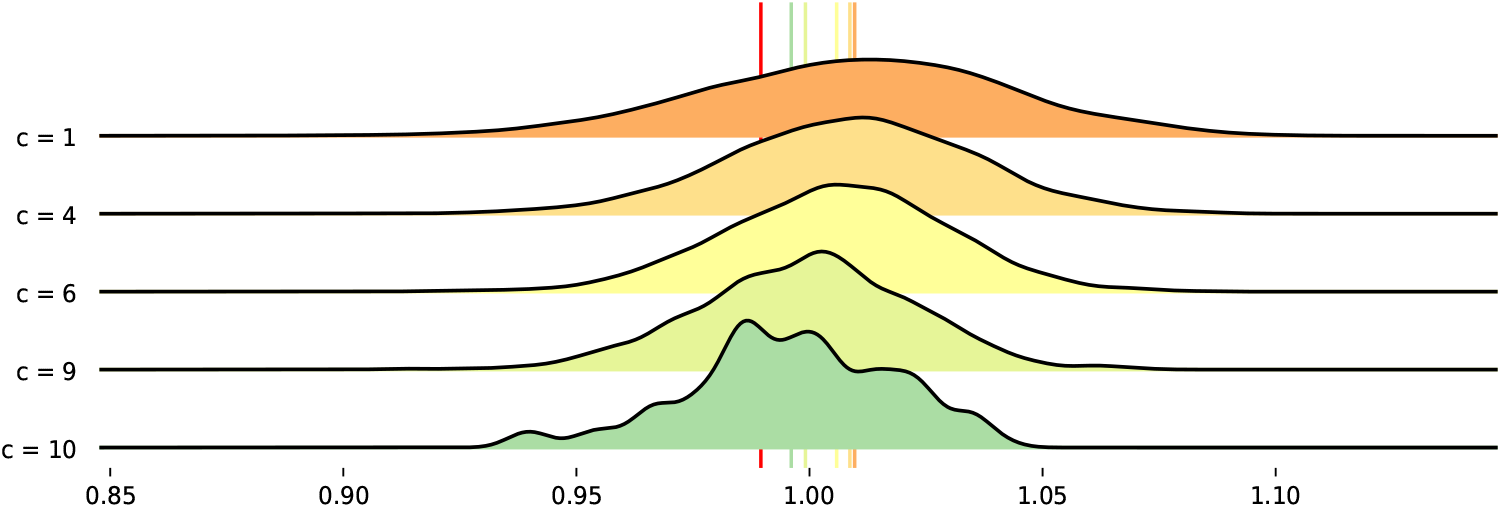
Predicted distribution of *λ* for each ABC-PMC iteration. Vertical lines indicate the true population growth rate, *λ*, (in red) and corresponding predicted mean for each iteration.

### 3.3 The real case study

A similar efficiency exploration about the real-world dataset can be found in Appendix C. The efficiency of the ABC-PMC is more pronounced in this real case study. The summary statistics for all particles are keeping reduced during iterations. Meanwhile, by the end, the true acceptance rate 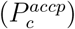 exceeds 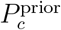 by over 50 times, and the total time for ABC-PMC amounted to only about 9% of the estimated implementation time for the single-iterated algorithm using just MCMC priors 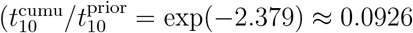 times).

The results concerning predictive performance are an extension of the initial exploration presented in Zhu et al. (2025). For a brief summary, we assessed the predictive performance of our models using IBM with ABC-PMC and MCMC samples, alongside GLMs to enable comparative analysis against traditional IPM approaches. The evaluation included both long-term and 1-step forecasts to examine demographic trends, using median values for predicted population sizes to mitigate outlier impacts.

Our findings indicated that GLM-based models generally yielded larger prediction mismatch (Figure 4) and less realistic simulations than GP-based models (Figure 5). In contrast, ABC samples improved accuracy greatly in GP models, providing more precise point estimates and relatively narrower 95% prediction intervals (PI) than the GP models without using ABC. Notably, according to NOAA (2013), the year 2012 holds the record for the warmest year in the historical record from 1895 to 2012 for the continental United States. We would not expect the proposed methods to effectively handle such extreme weather conditions.

**Figure 4:**
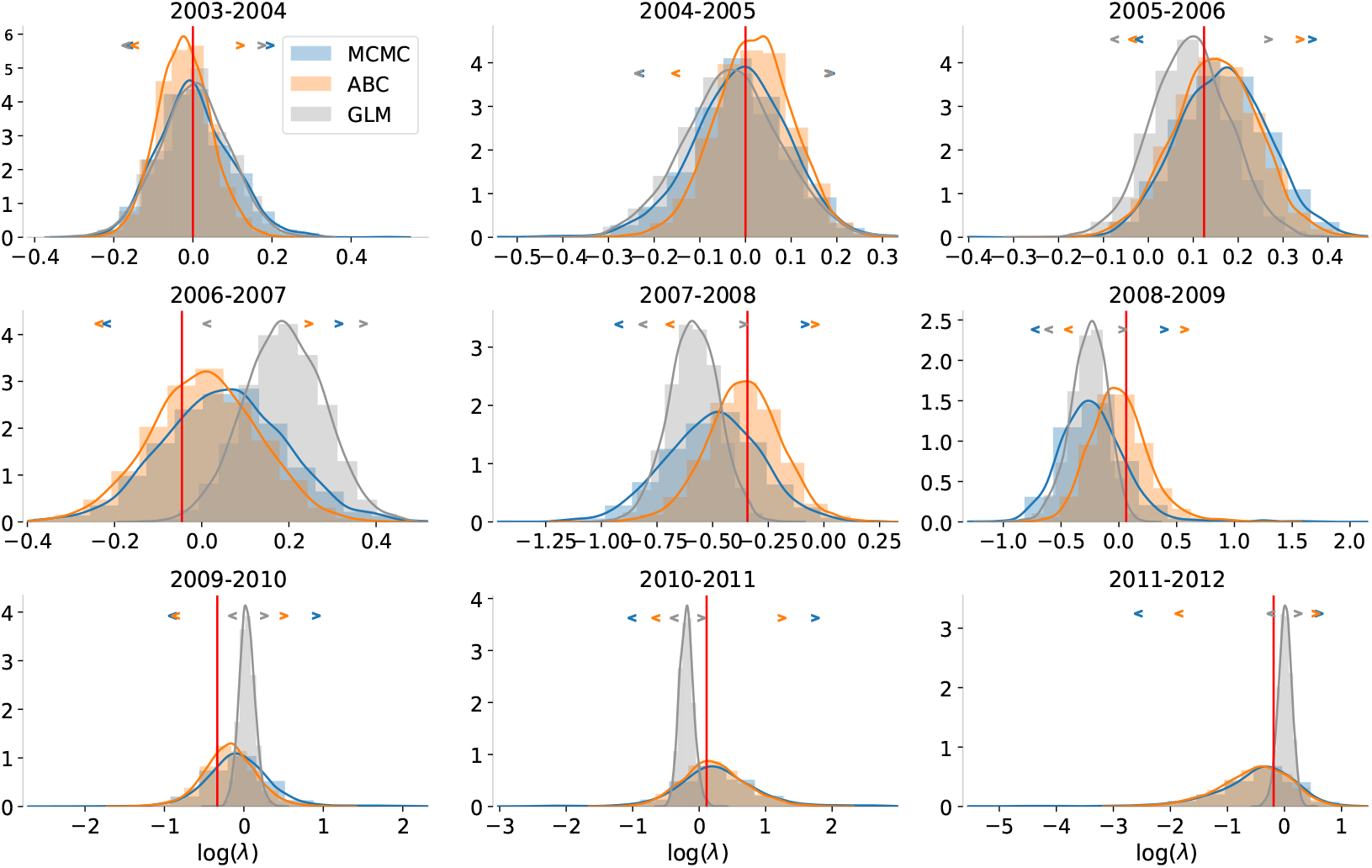
Histograms represent the density of logarithmic simulated population growth rates generated through **1-step** forecasting. The associated 95% prediction intervals are denoted by angle brackets, each in its corresponding colour, positioned at the top of each histogram. The red vertical line represents the true observed population growth rate.

**Figure 5:**
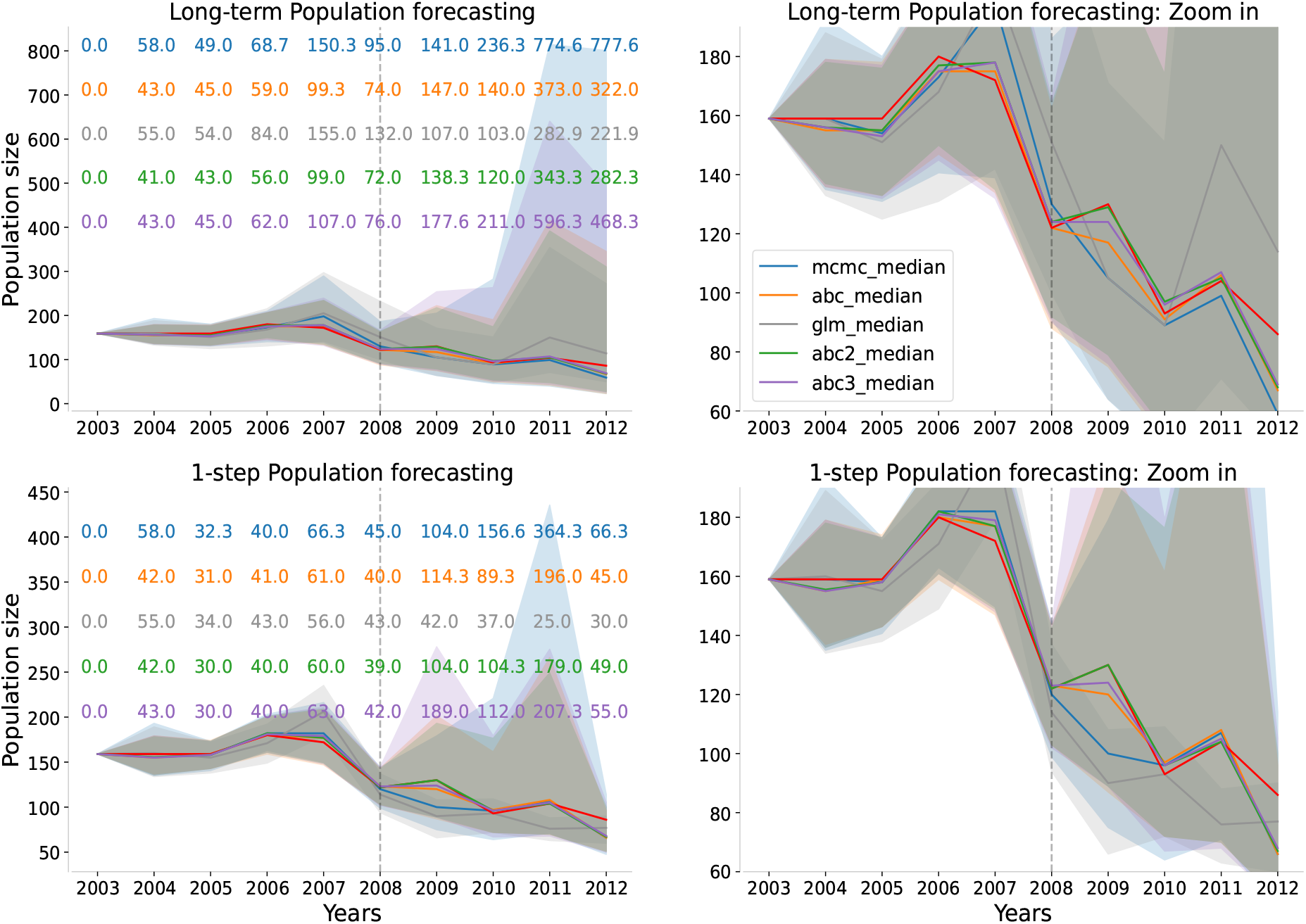
Two types of population size predictions, along with their corresponding 95% prediction intervals (shown as shaded areas), were generated using MCMC, ABC-PMC and GLM samples through 5,000 IBM simulations. The blue predictions were based on MCMC samples, while the orange and grey ones were based on ABC-PMC and GLM posterior samples, respectively. By separate replications, two other sets of ABC posterior samples generated through ABC-PMC are depicted here. These two additional simulations are also visually represented in green and purple. The widths of the corresponding 95% prediction intervals are indicated by their respective colours on the left-hand side of the figure. The point estimations are represented by solid lines in their corresponding colours. The red line represents the true observed population size.

The stochastic nature of the ABC algorithm, influenced by sampling randomness and adaptive threshold adjustments, leads to variability in posterior samples and corresponding PI, which can be narrower or wider. Figure 5 additionally presents forecasts from other two ABC-PMC implementations, illustrating varied uncertainties (shown as shadow areas) but consistently accurate point estimates (solid lines). Variations in PI generated by different replications of our proposed ABC-PMC algorithm could raise concerns about the convergence of ABC posterior samples. These concerns will be discussed in detail in section 4.

## 4 Discussion

This study adapts ABC into a viable inference framework for modular models, aiming to address a key limitation of the traditional “fit independently, then combine” approach. By applying ABC, our method enables the interactions among sub-models to be inferred from data rather than imposed by model assumptions. This makes ABC a practical option for high-dimensional modular models, even where sub-model interactions are important but challenging to specify explicitly.

To further improve ABC’s applicability, we developed a modified ABC-PMC sampler designed to better handle modular models, especially those incorporating machine learning-based sub-models such as GP models. Our method leverages pre-computed MCMC samples to refine both the prior and proposal distribution, greatly reducing rejection rates and computational cost. After a one-off preprocessing step (standard MCMC sampling for each sub-model), the time required to generate 10,000 proposals is reduced from over 20 days to under one minute, making ABC-PMC feasible for large-scale modular models.

We applied our method to an ecological case study using an IPM for *C. flava*. The results of the simulated and the real case studies demonstrate improved computational efficiency while preserving the quality of the inference. While the case study focuses on ecology, the method is applicable to a broad range of modular models where capturing interactions among sub-models is essential.

While our proposed ABC-PMC alleviates certain challenges of traditional ABC techniques, some limitations remain. For instance, constraining the proposals to precomputed MCMC samples restricts full exploration of the parameter space. We conclude with some final remarks on questions that remain unresolved.

### Did the ABC algorithm converge?

Figure 5 may raise concerns about the convergence of our ABC posterior samples. If the three sets of ABC posterior samples were obtained through independent replications, with the only difference being the replication itself, does the variability in their PIs suggest that our ABC-PMC algorithm have not fully converged? This is a valid concern, but achieving perfect convergence in ABC-PMC is inherently difficult.

The most straightforward reason is that, in practice, the ABC acceptance threshold cannot be arbitrarily small. Computational resources prevent us from enforcing stricter acceptance conditions while keeping the total runtime feasible. In our three ABC-PMC runs, the final thresholds reached approximately 9.104, 9.135, and 9.148, with acceptance rates around 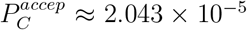. The total implementation time was 11761.89 minutes, approximately 8 days (see Table 2), making further reductions impractical. Notice that these threshold (9.104, 9.135, 9.148) are provided for reference but should not be directly comparable, as they were normalized using different MAD values.

Another factor is the design of the prior and perturbation kernel. Our prior restricts the parameter space to a finite set constructed by MCMC samples, which may not always contain values that satisfy a stricter threshold. The perturbation kernel, allowing only one sub-model to be perturbed at a time, further limits exploration. Since it may result in early elimination of potential good samples, reducing the diversity of available candidates in later iterations.

However, the final ABC posterior samples we obtained do not seem overly concerning and appear reasonably stable. As shown in Figure 10 in Appendix C, most hyper-parameters exhibit consistent marginal posterior distributions across runs, with only small mode shifts in `sur_ltmax` and `sur_ltmin`. In addition, the estimated population sizes from the three ABC repetitions show strong agreement, with overlapping point estimates (Figure 5).

While limited computational resources remain an objective constraint we cannot overcome, future work could explore ways to relax prior constraints, for broader parameter space exploration.

### Limitations of employing MCMC samples as ABC priors

With the nature of modular models, we decompose their hyper-parameters, ***θ*** *∈ 𝒫*, and locally store its distinct components. Our tailored ABC sampler then constrains the prior to a collection of points, denoted as *π*(***θ***) *∝ 𝒰* (***θ***) 𝟙 (***θ*** *∈* ***𝒳***), where the hyper-set ***𝒳*** *∈* ℝ^*dim*(***θ***)^ includes all possible combinations of unique groups of MCMC samples. This indicates that our method does not heavily rely on MCMC algorithms to provide reliable approximations of marginal distributions, but only needs them to return hyper-sets, representing the belief in a relative broad region where the target *π*_*ABC*_(***θ***|***s***^*obs*^) is likely to be found. Then, these necessary hyper-sets can be directly accessed by the proposed ABC-PMC, eliminating additional computation and processing time.

However, this convenience comes at a cost. Since the proposed ABC-PMC restricts sampling to ***𝒳***, a discretized subset of *𝒫*, it inherently assumes that high-probability posterior regions are well-represented within the precomputed MCMC samples. If key posterior regions are missing from ***𝒳***, they will not be explored by the proposed ABC-PMC.

A natural way to alleviate that is to increase the number of MCMC samples. However, this introduces additional storage and computational costs without necessarily resolving the fundamental limitation, especially when the MCMC posterior shape differs considerably from the shape of the ABC target. To illustrate, if the marginal ABC mode for a sub-model resides in the low MCMC posterior area, the pre-stored MCMC samples in this low probability region will be sparse. Consequently, a great number of ABC samples could be heavily concentrated on a single or few MCMC samples (from a marginal perspective), resulting in a very low effective sample size and reduced uniqueness for that sub-model. In our study, for instance, only 37 out of 200 samples were unique for the vital rate `fec`.

One possible approach to mitigate this limitation is to deliberately relax the sub-model posterior during MCMC sampling. This aims to produce more diffuse samples and construct a broader hyper-set ***𝒳***. Specifically, for a sub-model with parameters *θ*_*v*_ and data *y*_*v*_, instead of sampling from the standard posterior *π*(*θ*_*v*_)*p*(*y*_*v*_ | *θ*_*v*_), one could use a tempered version:

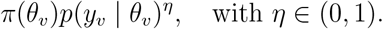

This approach, sometimes referred to as *tempered MCMC* or *power posteriors*, produces more diffuse samples (Friel & Pettitt (2008); Neal (1996)). The resulting marginal samples cover a wider region in parameter space, potentially improving the overlap between ***𝒳*** and the target ABC posterior *π*_*ABC*_(***θ***|***s***^*obs*^).

However, this comes with two costs: it reduces the influence of the data *y*_*v*_, so the resulting distributions no longer match the standard posterior implied by the model; and it introduces an additional tuning parameter, *η*. In our case, since these results are only used as priors for the ABC stage, the first cost may be acceptable. As for the second, its impact depends on how the method is implemented in practice.

## Acknowledgements

For the purpose of Open Access, the author has applied a CC BY public copyright licence to any Author Accepted Manuscript version arising from this submission.

## Summary statistics in ABC algorithms

Together with the choice of the sampling method, the selection of summary statistics has long been recognised as critical to the quality of ABC results. They all have a direct impact on the quality of the ABC approximation (Prangle (2017); Sisson et al. (2018)). ABC methods approximate the posterior distribution based on a set of assumptions, which include ensuring that the selected summary statistics ***s*** are representative (enough), the tolerance *h* is small (enough), then a good (enough) approximation of the posterior can be generated (Marin et al. (2012)).

Among the assumptions, the most tricky one is probably that we can expect that inferences about ***θ*** should involve the data only through the summary statistic ***s***. This assumption can be relaxed when the conditional density *π*(*y*, ***s***|***θ***) is independent or invariant of ***θ***, for which, we call the summary statistics ***s*** are (joint) sufficient for ***θ***. In this special case, there is no longer an ‘approximation’, while the outputs are exactly produced from the true posterior density, i.e. instead of *π*(***θ***|*y*) *≈ π*(***θ***|***s***), we have

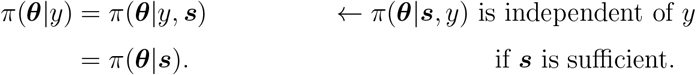

However, in a situation where the likelihood function is even unknown beyond a data generation procedure (otherwise, there would be no need for implementing ABC methods), finding low-dimensional sufficient statistics is a big challenge and could even be (nearly) impossible (Marin et al. (2012); Blum et al. (2013); Sisson et al. (2018)). Not to mention, we are still not sure if such low-dimensional sufficient statistics exist in the context of ABC.

According to the Pitman–Koopman–Darmois theorem, for independent identically distributed data points from a model *not* belonging to the exponential family, the dimension of any sufficient statistic remains unbounded and keeps increasing with the sample size. Given that ABC methods are generally not required for models from the exponential family (since their likelihoods are usually not intractable), the theorem strongly suggests that the low-dimensional sufficient statistics we are looking for are not likely to exist (Prangle (2018)). As using sufficient statistics is often not very likely in real-life situations, the choice of summary statistics play an even more important role in this case. It is directly responsible for the quality of the posterior approximation in ABC scenarios, based on the assumption *π*(***θ***|*y*) *≈ π*(***θ***|***s***). Consequently, methods for systematically selecting low-dimensional insufficient summaries are necessarily required.

In the absence of sufficient statistics in practice, it is intuitive to increase the number of summary statistics employed in the ABC samplers (to increase the amount of information available and thereby reduce the inference bias due to the irrevocable loss of information). More formally, relying on the rough idea of sufficiency, we may continue to add summary statistics and run ABC until the resulting posterior approximation stabilises (which is called Approximate sufficiency in Joyce & Marjoram (2008)). However, it is not always feasible to keep adding more and more summary statistics to the ABC samplers. The most apparent drawback of this approach is that it may suffer from the curse of dimensionality. The main issue is that, even with the correct model and the true parameter values, high-dimensional simulations rarely match fixed observations well due to numerous random components that must match the target simultaneously (Prangle et al. (2018)). This is why selecting appropriate summary statistics still remains one of the most challenging aspects of implementing ABC in practice. It is not an easy task to balance the need for a large number of summary statistics — to capture most of the information about the observed dataset — with the desire to keep the dimensionality low — to minimise random discrepancies between the summary statistics of the observed and simulated data.

Another potential risk of keeping adding more summary statistics with brute force is that it may introduce additional noise, leading to biased inference results (Beaumont et al. (2002); Csilléry et al. (2010); Sisson et al. (2018)). The toy example considered in Sisson et al. (2018) illustrated the issue.

### A toy example

The observed data *{y*_*i*_*}*_1⩽*i*⩽*n*_ were assumed to follow a Poisson distribution, *y*_*i*_ *∼ Poi*(*λ*), with parameter *λ ∼* Γ(*α, β*). This setting results in a closed-form posterior distribution for *λ, λ*|*y*_1_, …, 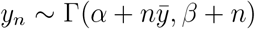, where 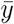 is the sample mean. Although the sample mean is a sufficient statistic for *λ*, they also considered using the sample standard deviation *v* as an additional (insufficient) summary statistic to estimate *λ* using ABC, given that the mean and variance of a Poisson distribution are both equal to *λ*. Given the parameters *α* = *β* = 1 and observations *{y*_*i*_*}*_1⩽*i*⩽5_ = (0, 0, 0, 0, 5), the ABC approximations are shown in Figure 6. Notice that, this setting is, in general, not going to happen in the real practice with ABC. There is no need to require ABC when likelihood is tractable.

**Figure 6:**
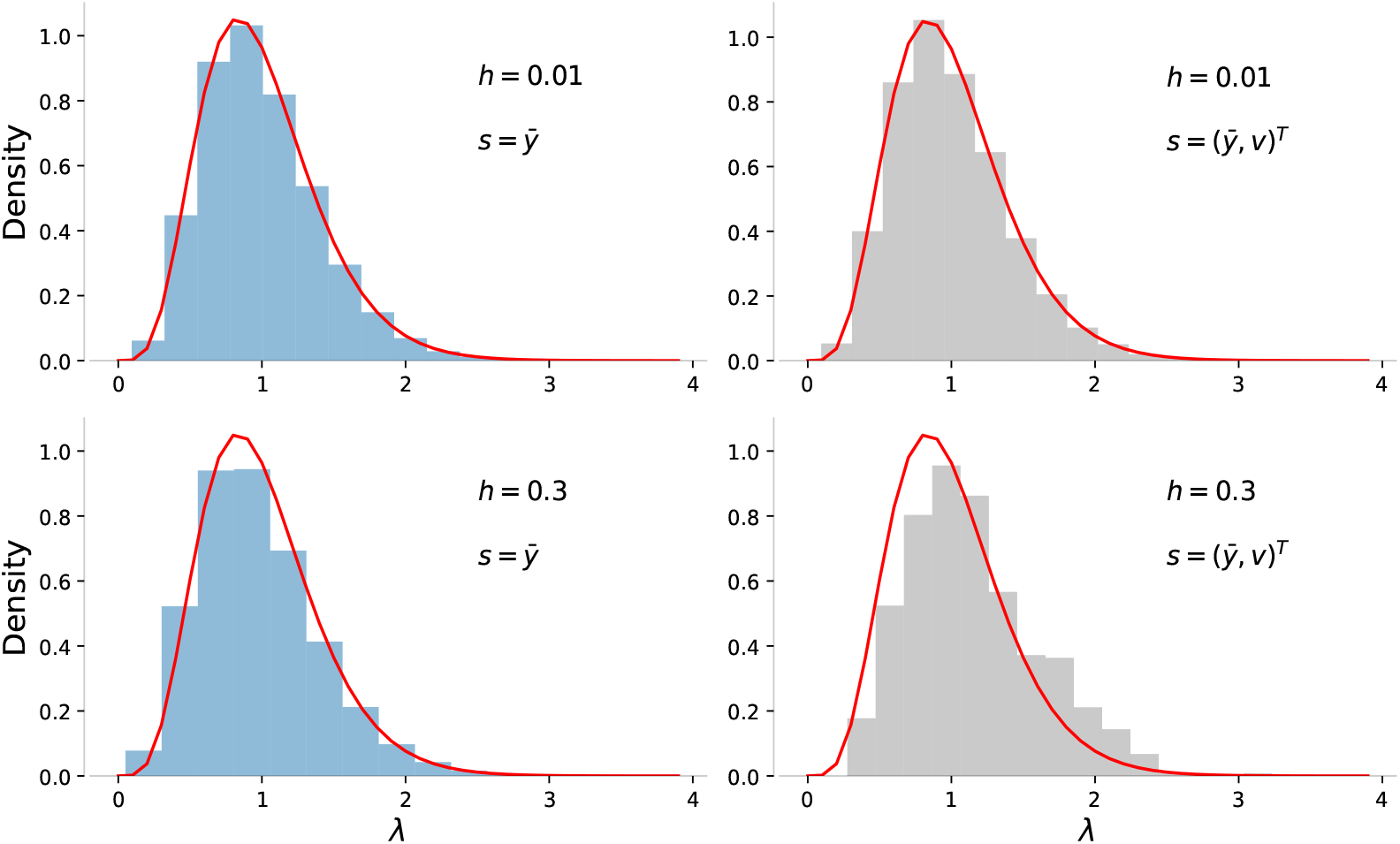
Density histograms of ABC posterior approximations. The approximations were generated through ABC Rejection samplers with the Euclidean distance metric and two different bandwidth values. That is, with *h* = 0.01 and *h* = 0.3, the acceptance probability *𝒦* in Equation 1 was defined as 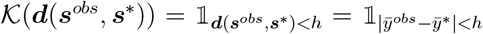 for plots on the left column, and 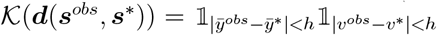 for plots on the right column. The red curves indicate the true posterior distribution Γ(6, 6).

When the sufficient statistic 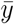 was solely employed for the ABC, the approximation near-perfectly matched the true posterior under a very narrow bandwidth value of *h* = 0.01. However, when the bandwidth got slightly wider *h* = 0.3, a minor fat-tailed pattern emerged in the approximation, deviating slightly from the true posterior distribution Γ(6, 6). The difference, while minor, emphasised the fact that the set of *λ* capable of reproducing 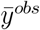 within a non-zero tolerance is (possibly similar, but still) different from the set that can exactly reproduce 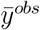. On the other hand, when the ABC approximation was jointly determined by sample mean and sample standard deviation, a strong restriction of *h* = 0.01 made the effect of the new summary statistics *v* almost negligible, leading to a similar approximation as using the sample mean only. However, when the ABC algorithm allowed for a more generous matching with *h* = 0.3, insufficient statistics *v* had additional opportunities to contribute more to the ABC approximation. The extra information provided by *v*^*obs*^ acted as ‘noise’, pushing the resulting posterior approximation away from the true posterior and producing a biased inference result. Thus, we should not expect that ABC methods based on the idea of approximate sufficiency would provide us reliable approximations (Sisson et al. (2018)), or at least not always.

Let us delve deeper into this example to see if we can gain any additional insights into ABC applications. The sample data *{y*_*i*_*}*_1⩽*i*⩽5_ = (0, 0, 0, 0, 5) yields a sample mean of 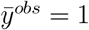 and a sample standard deviation of 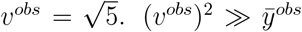 indicates an over-dispersed pattern, which is inconsistent with the common observations generated through a Poisson model. When both the sample mean and sample standard deviation are employed in ABC, increasing the threshold *h* results in more simulated parameters *λ*\* stand for larger values due to the term 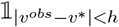 in 𝒦 (***d***(***s***^*obs*^, ***s***\*)), meanwhile, there are still quite a part speak for small values due to 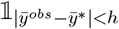. Then, it is not surprising to obtain a such ABC approximation that was skewed towards larger values when *h* = 0.01 became to *h* = 0.3 (the histograms on the right column in Figure 6).

We now gain a better understanding of the ‘noise’ generated by the summary statistic *v*. In this example, given the true model and the corresponding parameter, a set of observations containing information that was ‘inconsistent’ with the true model was generated. At that moment, an insufficient summary statistics bringing that piece of inconsistent information was employed into ABC methods, which, therefore, produced inference results that were not consistent with the actual posterior, namely the biased results. Now, the question is whether we can devise a strategy to reduce the impact of such summary statistics containing inconsistent information, and then obtain less biased inference results?

However, unlike in this given example, identifying which summary statistics are likely to bringing inconsistent information is already a big challenge when addressing real-life problems, let alone trying to reduce their impact. The task requires the knowledge of the true model behind, which might never be obtainable in the common practice. In this case, all we can do is, as with other approaches doing statistical inference, assume that the model is correct and make inference only about the parameter values. Under this assumption, summary statistics which are more intend to be unusual can be detected by simulating datasets from the given ‘true’ model. Then, relatively small weights could be assigned to these statistics to mitigate the impact of inconsistent information in the resulting ABC approximations.

Figure 7 displays the posterior approximation generated through an ABC rejection sampler that utilises a Euclidean metric-based weighting strategy, which has been commonly employed in previous studies, such as Hamilton et al. (2005); Luciani et al. (2009). The weights *w*_1_ = 1*/*0.8 and *w*_2_ *≈* 1*/*1.604 were defined as the inverse of median absolute deviations for 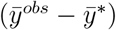 and (*v*^*obs*^ − *v*\*) respectively (Csilléry et al. (2012)). By giving a higher weight to 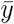, the new ABC posterior approximation was pulled back from the higher values (Figure 7 right). Although the impact of *v* was not completely eliminated (Figure 7 left), we did not observe a clear difference between the estimated and actual mode. The approach employed here automatically evaluates the stability or stiffness of two different types of the “difference” (i.e. the summary statistics) between the simulated and observed data and assigns weights accordingly.

**Figure 7:**
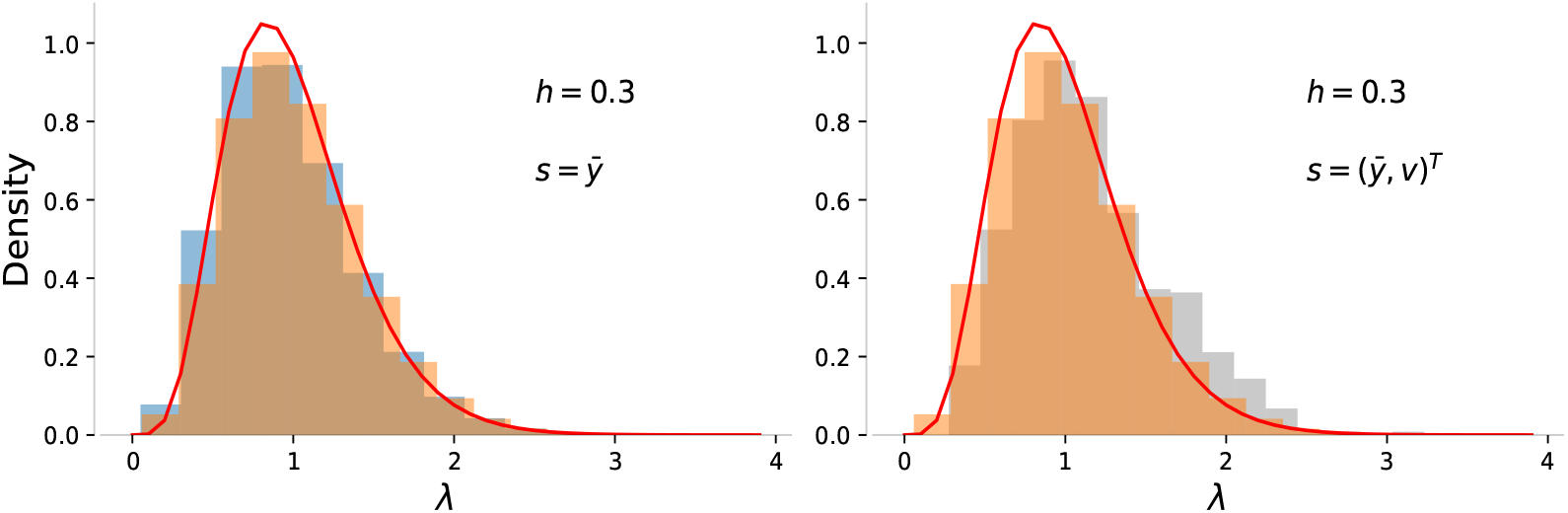
Density histograms of ABC posterior approximations. These two histograms are identical to the plots on the bottom of Figure 6. The additional orange shadow areas are the ABC posterior approximations based on the weighting strategy, where the acceptance kernel is defined as 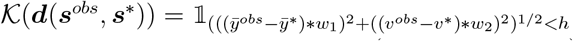 with *h* = ((0.3 *\* w*_1_)^2^ + (0.3 *\* w*_2_)^2^)^1*/*2^ to make the histograms comparable (from some sense).

This weighting strategy could not be useful for non-informative summary statistics, as they may be ‘steadily’ far from the observed values, but based on the assumption that all summary statistics employed are sensitive to the parameters, this strategy makes intuitive sense. On the other hand, by viewing the weights as the inverse of some factors, this approach can be also considered as a normalisation method. In this case, the method uses normalisation factors to bring all the individual ‘differences’ to the same scale, i.e. 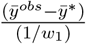 and 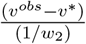 in this example, and to make them comparable for later use. We will discuss the weighting strategy in more details in Appendix B. In that section, following Prangle (2017), we will also discuss how the strategy can be developed to adaptive update their weights in each iteration of ABC-PMC and applied to increase the efficiency of ABC-PMC in our study.

There has been much progress regarding summary statistics in ABC scenarios. Some provide strategies to select or construct summary statistics that strike a balance between dimensionality and informativeness (Joyce & Marjoram (2008); Wegmann et al. (2009); Drovandi et al. (2011); Fearnhead & Prangle (2012); Scranton et al. (2014)); some concentrate on assigning appropriate weights for different statistics when they are under different scales or not equally informative (Beaumont et al. (2002); Luciani et al. (2009); Prangle (2017); Harrison & Baker (2020); Schälte & Hasenauer (2022)); some even bypass summary statistics altogether and compare complete datasets directly (Bernton et al. (2017); Gutmann et al. (2018); Bernton et al. (2019)). We refer Blum et al. (2013) and Prangle (2018) which provide detailed reviews of the ABC summary statistics.

### B Explanations for the ABC-PMC Modifications

This section describes the motivations and thought process behind the three modifications made to our developed ABC-PMC. We aim to share why and how we decided to make these adjustments in our case, which may be helpful for others looking to apply or develop these methods in their studies.

1. We utilise a tailor-designed prior distribution *π*(***θ***) and perturbation kernel *𝒫* (*·*|*·*) in our implementation of ABC-PMC to better suit our specific case of modular models with highly flexible sub-models, like GP models.

#### The problem

The practical application of GP models has long been hindered by the computational complexity involved (Cheng & Boots (2017); Schulz et al. (2018)). In our case of implementing ABC, obtaining a value for GP hyperparameters is (nearly) cost-free since a value can be simply drawn from the proposal distribution rather than being estimated. However, simulating populations using those specified hyperparameter values remains very expensive. The computational complexity is dominated by computing the GP predictive distributions, which require a complexity of *𝒪*(*N* ^3^) when solving or approximating inference with *N* training data points through the Cholesky decomposition.

In our experiments of IPMs, the entire process — manually feeding an assigned value of ***θ*** into GP models of the five vital rates (processed in serial) and preparing them ready for the simulations — took about 3 minutes on average on a machine equipped with the 10-core Apple M1 Max processor. In this way, it would take at least 20 days to achieve 10,000 attempts. It is worth mentioning that in most ABC-PMC applications, 10,000 *acceptances* are not sufficient for one iteration, let alone for just 10,000 *attempts*.

The situation can become even worse when ***θ*** is of high dimension in modular models, as interactions between sub-models can amplify the effects of any inconsistency. Considerable workloads on computing resources can be exerted by the choice of inefficient perturbation kernels during the sampling process (Filippi et al. (2013)). In our scenario, with the commonly adopted multivariate normal perturbation kernel (e.g. Beaumont et al. (2009)), a dramatically high rejection rate was observed. The proposal movements, even if slightly inconsistent or inappropriate, could generate very unrealistic vital rates, resulting in simulated populations that often did not resemble the observations. The acceptance rate was less than 0.1% — this was the case even when the movements were proposed based on a population of very promising particles and with a very gentle acceptance threshold determined by background knowledge.

In a high-dimensional problem, designing a perturbation kernel that can efficiently move across all dimensions of ***θ*** simultaneously is challenging (Everitt & Rowińska (2021)). Given such a high rejection rate, one can imagine the immense computational time required as the process of computing approximations in GP models is repeated millions and billions of times in the conventional ABC-PMC. Even with parallel computing on a powerful 64-core server, completing the ABC model fitting within a reasonable time frame may still be unfeasible.

#### The solution

Recall that, in modular models, the most commonly adopted model fitting process comprises two key steps. Firstly, each sub-model is independently parameterized from the others. Secondly, these multiple pieces of individual models are combined to construct a comprehensive model. This procedure reminds us of a major distinction between the modular models and some other complicated models with analytically intractable likelihoods. Being a modular model, it can therefore also be decomposed back into those simpler baseline models, each pertaining to individual sub-model. For those sub-models, particularly the GP models in our study, a plenty of robust tools have been developed to efficiently draw samples from the respective individual target posterior. Why not follow the initial step of the standard modular model fitting procedure to gain a rough idea about the posterior distribution of the parameters, for example, by obtaining MCMC samples? Then, we can re-use these MCMC samples for each individual vital rate (serving as our ‘background knowledge’ about ***θ***) to construct a more efficient ABC-PMC for exploring their joint behaviours.

In other words, instead of using non-informative priors, like a high-dimensional uniform distribution with wide boundaries *𝒰* (***θ***), we can specify our prior for the hyperparameter vector ***θ*** to be confined by a set of points *π*(***θ***) *∝ 𝒰* (***θ***) *𝟙* (***θ*** *∈* ***𝒳***). Here, the hyperset ***𝒳*** *∈* ℝ^*dim*(***θ***)^ includes all combinations of different groups of MCMC samples and represents our belief in a broad region where the target *π*_*ABC*_(***θ***|***s***^*obs*^) is likely to be found. After obtaining, for example, *M* different MCMC samples for each of *V* individual submodels, hyper-set ***𝒳*** contains *M*^*V*^ possible points.

To avoid a dramatically high rejection rate resulting from inappropriate proposals, it is also necessary to re-design the perturbation kernel *𝒫* (*·*|*·*). Generally, the perturbation kernel in ABC serves two primary purposes: (i) guiding particles towards regions of high posterior probability, and (ii) enhancing particle diversity (thereby facilitating a more extensive exploration of the parameter space) (Filippi et al. (2013); Sisson & Fan (2018)). In our special case, we also require a perturbation kernel from which new particles can be easily and readily sampled, and, meanwhile, computing the transition density between any two particles should not impose great additional computational load.

Drawing upon the MCMC samples, we formulate a straightforward component-wise perturbation kernel *𝒫* (*·*|*·*). Given a particle accepted in the previous iteration, 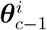, a new movement is proposed by randomly selecting one of the *V* sub-models and replacing all corresponding parameters with a randomly chosen sample from the *M* MCMC samples (Line 15 in Algorithm 3). Accordingly, the resulted proposal distribution becomes to 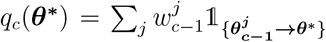 for iteration *c* = 1, …, *C*, where 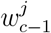 is the corresponding weight for particle 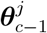 accepted in the previous iteration. 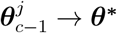 is a statement about whether it is possible to transit from point 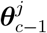 to point ***θ*\*** through the perturbation kernel (Line 21 in Algorithm 3).

In each perturbation, the kernel *𝒫* (*·*|*·*) is designed to make only small adjustments to increase the acceptance probability. It can explore around the current particles more thoroughly, accepting only those new particles that can produce a simulation similar to the observed one. After obtaining the required number of acceptances, the proposal distribution *q*_*c*_ (in Equation 3) is automatically updated by incorporating these more promising particles, so that, it will be able to propose values that are more likely to lie in high posterior regions for the next iteration. Through this iterative process, *q*_*c*_ based on the proposed *𝒫* (*·*|*·*) is expected to continually improve and to ultimately evolve into a highly efficient proposal distribution for the final round.

When using ABC-PMC with this configuration, the computational overhead associated with the approximations in GP models is no longer a big concern. The parameters for these approximations have already been calculated during MCMC simulations, so all that is needed is additional storage space to cache the MCMC results locally and retrieve them as needed. Similarly, for the Cholesky decomposition required to compute predictive distributions, due to the use of a fixed set of MCMC samples for each sub-model, predictions with the same MCMC sample occur much more frequently. In such cases, pre-doing the calculations that are independent of the test points and reusing them between predictions can substantially alleviate the computational burden. Additionally, there is also no need for additional resources to calculate transition probabilities between particles, as they will be a constant of 1*/*(*V M*).

2. As a toy example demonstrated in Appendix A, **an adaptive weighting strategy** is employed to combine selected summary statistics into a single ‘universal’ distance, measuring the difference between observed and simulated datasets.

#### The problem

In ABC, several distinct summary statistics should be chosen to represent various aspects of the dataset: if only one type of the summary statistics is employed in ABC-PMC, the sampler may be excellent for estimating the parameters for a particular aspect, but can be very poor and less-constrained for the others (Hamilton et al. (2005)). Subsequently, for parameterising multiple sub-models simultaneously, incorporating all the selected summary statistics as a multi-dimensional vector into the ABC sampler is essential, ensuring a comprehensive representation of the observed dataset.

Typically, ABC samplers use a “rectangular” acceptance region when dealing with multi-dimensional summary statistics (Pritchard et al. (1999); Scranton et al. (2014)), as the indicator function in Equation 3. Thus, a parameter is accepted only if all summary statistics satisfy their respective thresholds simultaneously.

The method assumes equal contributions from all summary statistics to the estimation of the target posterior when accepted. However, when summary statistics are constrained to be interpretable in reality, they may carry overlapping information, causing some to be less informative in the presence of others. This complicates the selection of appropriate thresholds 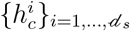 to prevent less-informative statistics from over-affecting the parameter estimations and ensure that all statistics contribute equally.

Additionally, applying a such “rectangular” acceptance region in our study may suffer from the curse of dimensionality. Despite our efforts to keep the number of summary statistics as small as possible, having a dimension such as *d*_*s*_ = 25 is still quite large (one summary statistic per vital rate over five years). With such a high-dimensional vector of summary statistics, the ABC algorithm inevitably faces a very high rejection rate in the “rectangular” acceptance region, as it only accepts particles that meet all conditions simultaneously. These challenges further complicate the estimation of parameter values.

To alleviate the computational overhead due to the high rejection probability in the “rectangle”, Beaumont et al. (2002) combined the summary statistics through weighted *L*^2^ norm, 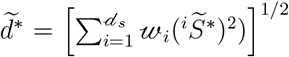, transforming the region into a “sphere” to increase the acceptance rate for the samples. To make the normalisation in a more appropriate way, the weight 𝓌_*i*_ of the *i*th summary statistic is usually taken as the inverse of its estimated dispersion measure calculated from the prior predictive distribution, such as the empirical standard deviation (Beaumont et al. (2002)), for other examples, see Hamilton et al. (2005); Luciani et al. (2009); Erhardt & Sisson (2016); Prangle (2017); Rodrigues et al. (2018).

During our studies on IPMs, we found that certain hyperparameter combinations could yield biologically unrealistic vital rates, like an extremely high survival rate for the most reproductive group (which is not typically observed in reality). This discrepancy resulted in huge divergence between simulated and the observed populations. To address this issue, we normalise the summary statistics using the median absolute deviation (MAD) as suggested by Csilléry et al. (2012). Normalising through MADs is less sensitive to those large discrepancies (i.e. outliers) and can also intuitively reduce the impact of the summary statistics that are running on different scales, tending to stop the acceptance rules being dominated by the most variable summary statistic (Prangle (2017); Schälte & Hasenauer (2022)).

However, given the iterative nature of ABC-PMC, it appears inappropriate to employ the standard scaling strategy of utilising consistent MADs (which are estimated from the *initial* prior predictive distribution) throughout the *entire* ABC-PMC process. This is because the objective function will actually keep changing in each iteration, when, for example, a decreasing sequence of thresholds *{h*_*c*_*}*_*c*=0, …, *C*_ is employed. In this case, later rounds of ABC-PMC may yield realisations originating from distributions vastly different from the initial prior predictive distribution, making the normalising strategy meaningless. To prevent this problem in our version of ABC-PMC, we adopt an adaptive strategy to update the MADs in each iteration according to Prangle (2017).

#### The solution

The idea behind this modification is straightforward: updating the MADs based on simulations generated in the previous iteration to achieve more sensible normalisation in each round. Specifically, in Algorithm 3, we incorporate all particles in the previous iteration (including those rejected), to calculate the MADs. The reason for including rejected particles is to prevent bias when the target distributions between consecutive iterations are very different. If we rely only on accepted particles, the MADs might primarily reflect the criteria used in the previous iteration, which could no longer match the current target distribution. Including all particles would instead be less restrictive and can provide a more comprehensive picture of the samples generated based on the intermediate proposals. This helps MADs to be more representative of the broader sample set and not overly influenced by past criteria (Prangle (2017)).

Note that this modification may still be inadequate if the targets distributions in two successive iterations differ dramatically. In such cases, an alternative approach proposed by Prangle (2017) could be considered. Generally speaking, this alternation involves treating the accepted particles in Algorithm 3 as ‘inter-mediate’ particles, and using them to estimate the MADs for normalizing summary statistics generated by an additional layer of simulations. However, this approach requires a large enough set of ‘inter-mediate’ particles to produce reliable MAD estimations, which can impose a substantial computational burden.

In our study, the rejection rates were already extremely high in the latter rounds of Algorithm 3 (see Table 3). With a fixed computation budget, the additional cost required by introducing ‘inter-mediate’ particles prevented us from using small values in acceptance thresholds *{h*_*c*_*}*_*c*=0, …, *C*_. This might be the reason why this method performed even worse than Algorithm 3 in our experiment (not shown). Ultimately, we chose to stick with Algorithm 3 for the latter work, as a trade-off for the limited computational resources.

In summary, the ‘universal’ distance between the observed and a simulated dataset at *c*th iteration is defined by

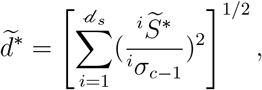

where 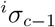 is the estimated median absolute deviation (MAD) (based on *c* − 1th generation of particles) for the *i*th summary statistic 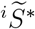.

**Table 3:** Results of applying ABC-PMC to GP-IPM for real case study. **Notations**: *δ*_**c**−**1**_: the quantile-based acceptance threshold used at step *c*, which was determined based on *α*_**c**−**1**_ quantile at step *c* 1 (so denoted by *δ*_*c*−1_). *δ*_−1_ and *α*_−1_ are for the initialisation. 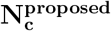: the proposed number of accepted particles required for the simulation step *c*. **N**_**c**_: the true number of accepted particles for the simulation step *c*. 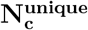: the number of unique particles among *N*_*c*_ acceptances. **n**(***𝒳***_**c**_): the total number of possible states of ***θ*** at simulation step *c*. Apart from **n**(***𝒳***_**0**_), the others were estimated through *n*(***𝒳***_*c*_)^*lb*^. 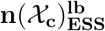: a lower bound for *n*(***𝒳***_*c*_) but estimated based of ESS. 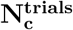: total number of trials to achieve the acceptance. 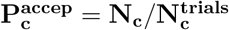: the acceptance rate for step *c*. 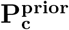: the acceptance rate for a single-iterated ABC-PMC, where the samples were drawn directly from the priors made up by MCMC samples. For the reliability, 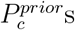 were computed based on 10,000,190 particles drawn from *π*(***θ***). To ensure comparability, 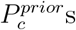 must be computed under the exactly same setting that produced 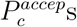. That is, for each *c* in *{*0, 1, …, *C}*, the 10,000,190 particles were normalised with MADs ***σ***_*c*−1_ first, the threshold *δ*_*c*−1_ then employed to compute 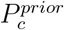. It took approximately 2,541 minutes to complete this single-iterated algorithm with 10,000,190 particles. **t**_**c**_: the total amount of time, measured in minutes, used to complete the *c*th iteration.

| | $c = 0$ | $c = 1$ | $c = 2$ | $c = 3$ | $c = 4$ | $c = 5$ |
| --- | --- | --- | --- | --- | --- | --- |
| $\alpha_{c-1}$ | 1 | 0.55 | 0.50 | 0.45 | 0.40 | 0.35 |
| $\delta_{c-1}$ | ∞ | 21.01 | 18.46 | 17.05 | 15.60 | 14.19 |
| $N_c^{proposed}$ | 400,000 | 240,000 | 200,000 | 140,000 | 100,000 | 60,000 |
| $N_c$ | 400,017 | 242,959 | 201,541 | 141,150 | 100,473 | 60,241 |
| $N_c^{unique}$ | 400,017 | 242,956 | 201,538 | 141,148 | 100,471 | 60,241 |
| $n(\mathcal{X}_c)$ | 5000<sup>5</sup> | $> 1 \times 10^{10}$ | $> 5.3 \times 10^9$ | $> 4.1 \times 10^9$ | $> 2.9 \times 10^9$ | $> 2.0 \times 10^9$ |
| $n(\mathcal{X}_c)_{ESS}^{lb}$ | 5000<sup>5</sup> | $> 1 \times 10^{10}$ | $> 5.3 \times 10^9$ | $> 4.1 \times 10^9$ | $> 2.8 \times 10^9$ | $> 1.9 \times 10^9$ |
| $N_c^{trials}$ | 400,017 | 440,000 | 670,000 | 890,000 | 1,270,000 | 1,710,000 |
| $P_c^{accep}$ | 1 | 0.5522 | 0.3008 | 0.1586 | 0.0791 | 0.0352 |
| $P_c^{prior}$ | 1 | 0.5510 | 0.2746 | 0.1215 | 0.0462 | 0.0152 |
| $t_c$ | 96.35 | 152.85 | 190.27 | 236.50 | 326.55 | 433.95 |

Table 3: Results of applying ABC-PMC to GP-IPM for real case study.
| | $c = 6$ | $c = 7$ | $c = 8$ | $c = 9$ | $c = 10$ |
| --- | --- | --- | --- | --- | --- |
| $\alpha_{c-1}$ | 0.30 | 0.20 | 0.15 | 0.10 | 0.088 |
| $\delta_{c-1}$ | 13.02 | 11.88 | 10.91 | 9.995 | 9.135 |
| $N_c^{proposed}$ | 30,000 | 20,000 | 10,000 | 1,600 | 200 |
| $N_c$ | 30,046 | 20,018 | 10,012 | 1,600 | 200 |
| $N_c^{unique}$ | 30,045 | 20,018 | 10,010 | 1,600 | 200 |
| $n(\mathcal{X}_c)$ | $> 1.2 \times 10^9$ | $> 6.3 \times 10^8$ | $> 3.8 \times 10^8$ | $> 1.9 \times 10^8$ | $> 4.5 \times 10^7$ |
| $n(\mathcal{X}_c)_{ESS}^{lb}$ | $> 1.1 \times 10^9$ | $> 5.6 \times 10^8$ | $> 3.3 \times 10^8$ | $> 1.5 \times 10^8$ | $> 2.6 \times 10^7$ |
| $N_c^{trials}$ | 2,180,000 | 5,390,000 | 11,900,000 | 11,410,000 | 9,790,000 |
| $P_c^{accep}$ | 0.0138 | 0.0037 | $8.413 \times 10^{-4}$ | $1.402 \times 10^{-4}$ | $2.043 \times 10^{-5}$ |
| $P_c^{prior}$ | 0.0041 | 0.0007 | $8.200 \times 10^{-5}$ | $7.700 \times 10^{-6}$ | $4.000 \times 10^{-7}$ |
| $t_c$ | 550.62 | 1362.4 | 3018.8 | 2903.7 | 2489.9 |

3. Additionally, the last modification concerns the quantile used to decide the **acceptance region**. Employing a constant quantile *α* in Algorithm 3 brings us a potential risk: the determined acceptance regions might not originate from subsets in a nested sequence converging to 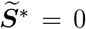 (Prangle (2017); Schälte et al. (2021)). To make particles from each successive iteration less diffuse, we explore various schedules and ultimately choose decreasing sequence of quantiles *{α*_*i*_*}*_0⩽*i*⩽*C*−1_ — to ensure that *δ*_*i*_ *< δ*_*i*+1_ for 0 ⩽ *i < C* − 1 and, therefore, expect that the populations of accepted particles gradually converge to the final target distribution in the limit (Toni et al. (2009)).

Under this quantile sequence, rejection rates could increase remarkably as *α*_*i*_ decreased. However, we only have a fixed computational budget that can be allocated at each simulation step. To balance computational time, we have to make an additional modification by gradually reducing the total number of accepted particles required (*{N*_*i*_*}*_0⩽*i*⩽*C*_). Under a reasonable computational time, we should turn up the *N*_0_ as large as possible. We then progressively reduce *N*_*i*_ for the subsequent iterations in a controlled manner to ensure that the decrease is not too rapid. This is important because a rapid decrease in *{N*_*i*_*}*_0⩽*i*⩽*C*_ would not provide enough time to gradually reduce *α*_*i*_, which is necessary to maintain a sufficient number of accepted perturbation (Marin et al. (2012)). By managing the reduction, we are able to obtain a final generation of accepted particles that is still of a reasonable size. In addition, we would also ensure that the decrease is not too slow. This is because, given the accepted particles in the previous iteration, the total number of possible states of ***θ*** is fixed under our perturbation kernel (see Table 2 for more details). A proper reduction in the number of acceptances allows us to propagate only the most promising particles, those with a high chance of 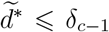 in step 19 of Algorithm 3, providing stability for the future samples.

### C Results for the real case study

**Figure 8:**
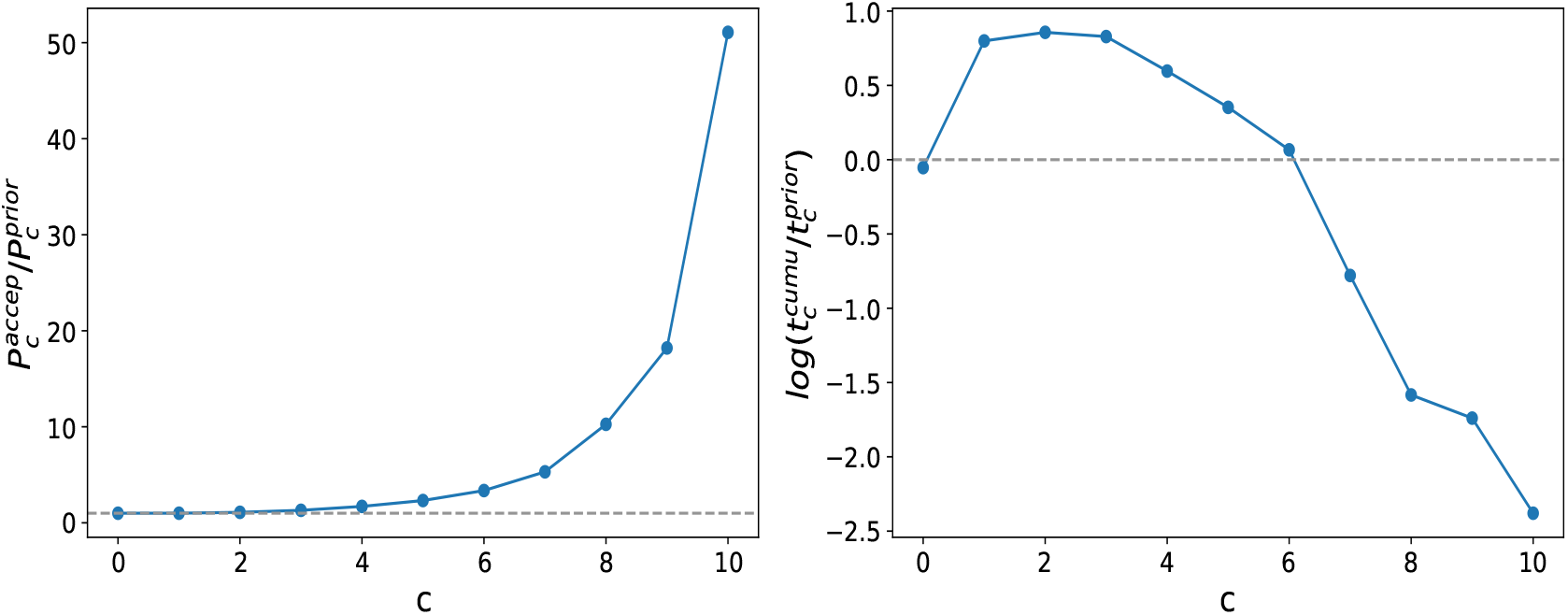
**Left:** The ratio of 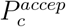 and 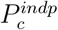 over simulation time *c*. The grey horizontal line cross the point representing 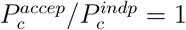. **Right:** The logarithmic ratio between the cumulative time spent on ABC-PMC, represented as 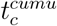, and the expected implementation time, estimated as 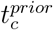. The 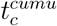 is obtained by adding up individual times for each single iteration *t*_*c*_ until the current iteration 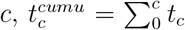. Instead of timing in real practices, 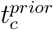 was estimated using the formula 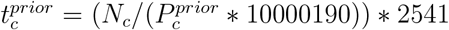 based on the 10,000,190 particles (see Table 3).

**Figure 9:**
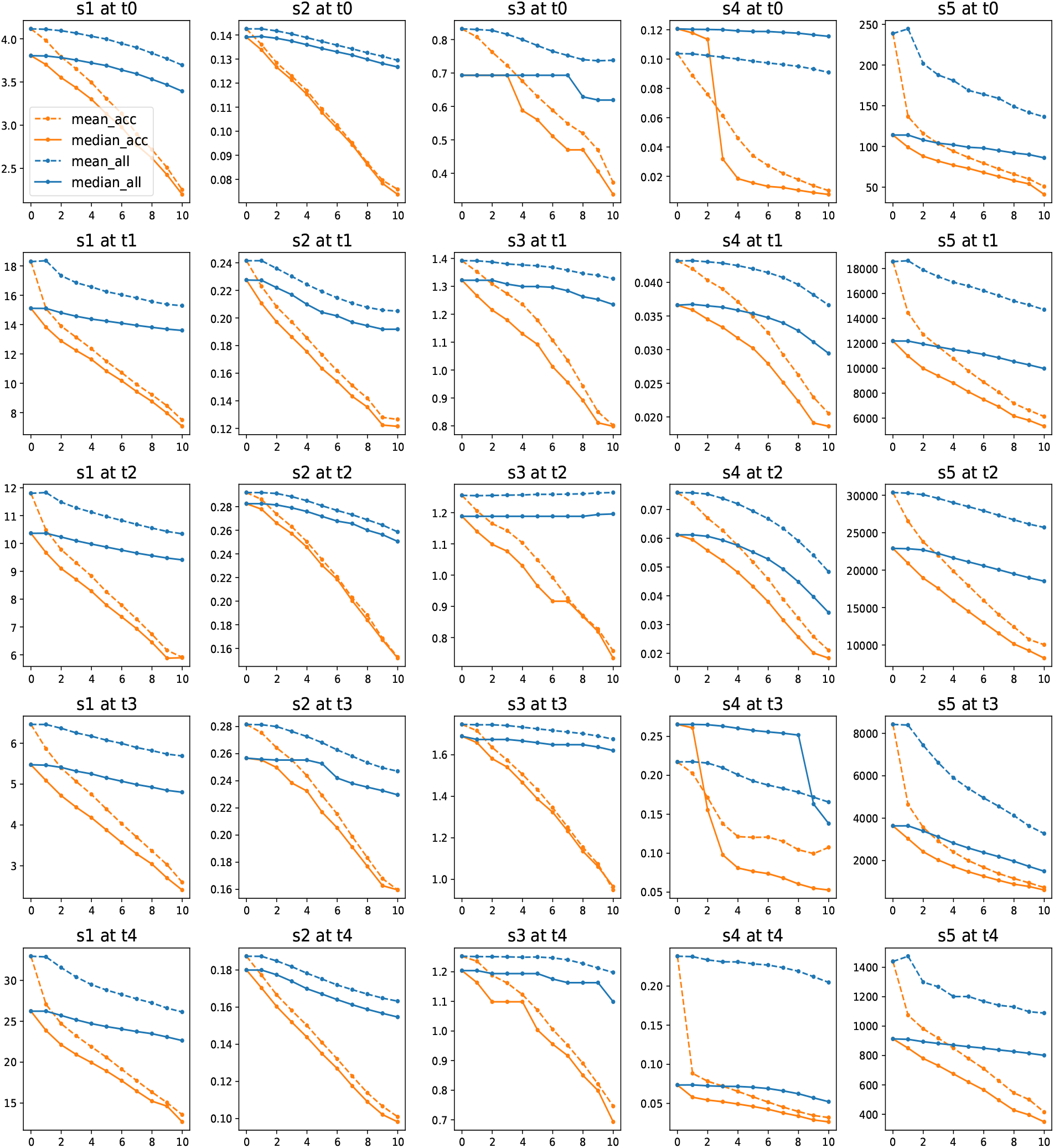
The mean (dashed line) and median (solid line) for the 25 summary statistics over the ABC-PMC simulation time *c* (*x* − axis). The orange lines describing the information of summary statistics for accepted particles only, while the blues stand for all particles including those rejected. **s1** to **s5** represent the five types of summary statistics, **t0, t1**, …, **t4** indicate year 2003-2004, 2004-2005, …, 2007-2008.

**Figure 10:**
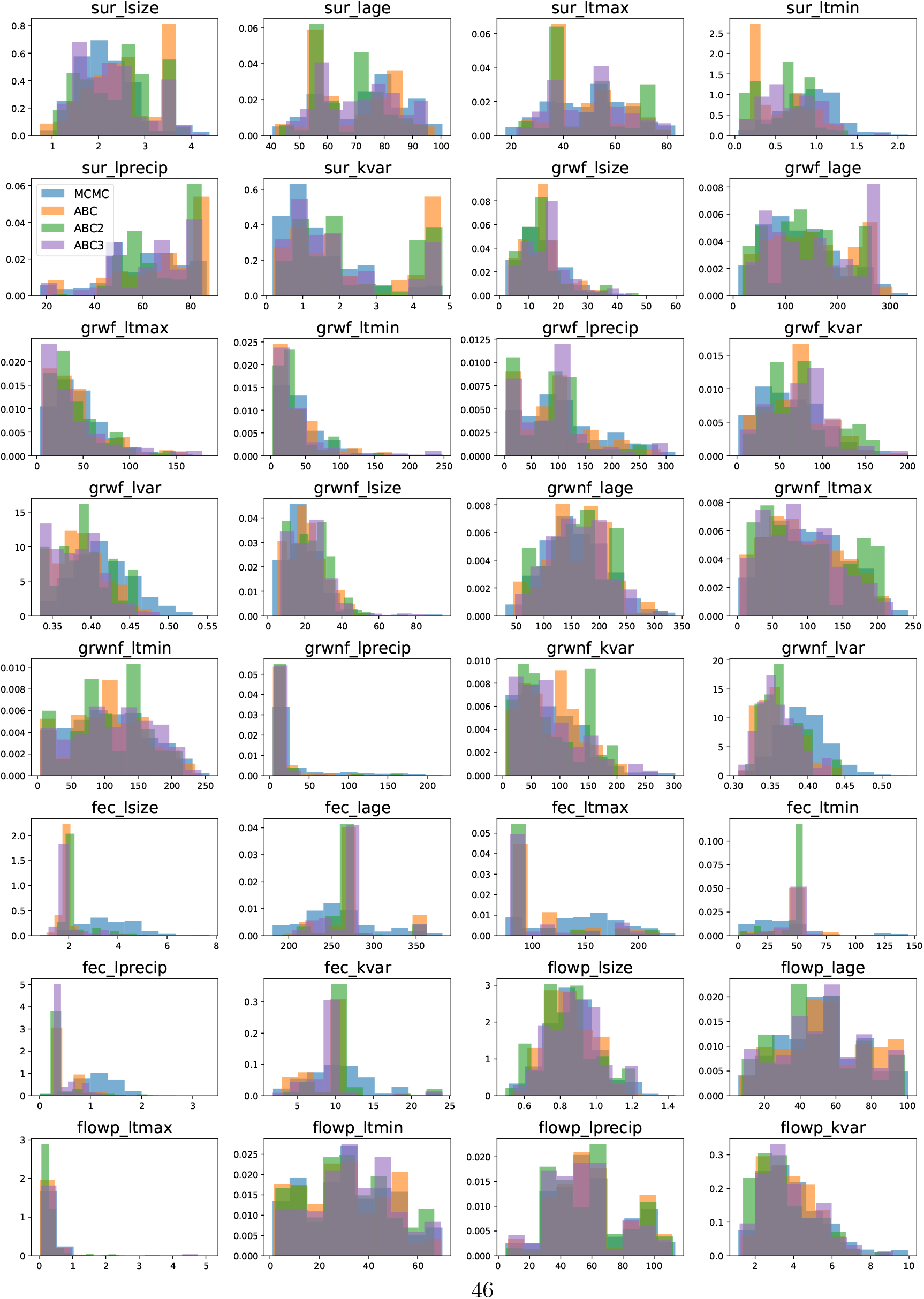
Density histograms of estimated marginal posterior distributions for all hyperparameters under GP and ABC_GP.

## References

Andrieu, C., Lee, A., & Vihola, M. (2018). Theoretical and methodological aspects of mcmc computations with noisy likelihoods. Chapman & Hall/CRC Handbooks of Modern Statistical Methods.

Barnes, C. P., Silk, D., Sheng, X., & Stumpf, M. P. (2011). Bayesian design of synthetic biological systems. Proceedings of the National Academy of Sciences, 108(37), 15190– 15195.

Beaumont, M. A., Cornuet, J.-M., Marin, J.-M., & Robert, C. P. (2009). Adaptive approximate bayesian computation. Biometrika, 96(4), 983–990.

Beaumont, M. A., Zhang, W., & Balding, D. J. (2002). Approximate bayesian computation in population genetics. Genetics, 162(4), 2025–2035.

Bernton, E., Jacob, P. E., Gerber, M., & Robert, C. P. (2017). Inference in generative models using the wasserstein distance. arXiv preprint 1701.05146, 1(8), 9.

Bernton, E., Jacob, P. E., Gerber, M., & Robert, C. P. (2019). Approximate bayesian computation with the wasserstein distance. arXiv preprint 1905.03747.

Blum, M. G., Nunes, M. A., Prangle, D., & Sisson, S. A. (2013). A comparative review of dimension reduction methods in approximate bayesian computation. Statistical Science, 28(2), 189–208.

Cappé, O., Guillin, A., Marin, J.-M., & Robert, C. P. (2004). Population monte carlo. Journal of Computational and Graphical Statistics, 13(4), 907–929.

Caswell, H. (2000). Matrix population models, volume 1. Sinauer Sunderland, MA, USA.

Chen, V., Yang, M., Cui, W., Kim, J. S., Talwalkar, A., & Ma, J. (2024). Applying interpretable machine learning in computational biology—pitfalls, recommendations and opportunities for new developments. Nature methods, 21(8), 1454–1461.

Cheng, C.-A. & Boots, B. (2017). Variational inference for gaussian process models with linear complexity. Advances in Neural Information Processing Systems, 30.

Csilléry, K., Blum, M. G., Gaggiotti, O. E., & François, O. (2010). Approximate bayesian computation (abc) in practice. Trends in ecology & evolution, 25(7), 410–418.

Csilléry, K., François, O., & Blum, M. G. (2012). abc: an r package for approximate bayesian computation (abc). Methods in ecology and evolution, 3(3), 475–479.

Drovandi, C. C. & Pettitt, A. N. (2011). Estimation of parameters for macroparasite population evolution using approximate bayesian computation. Biometrics, 67(1), 225–233.

Drovandi, C. C., Pettitt, A. N., & Faddy, M. J. (2011). Approximate bayesian computation using indirect inference. Journal of the Royal Statistical Society: Series C (Applied Statistics), 60(3), 317–337.

Ellner, S. P., Childs, D. Z., Rees, M., et al. (2016). Data-driven modelling of structured populations. A practical guide to the Integral Projection Model. Cham: Springer.

Ellner, S. P. & Rees, M. (2006). Integral projection models for species with complex demography. The American Naturalist, 167(3), 410–428.

Erhardt, R. & Sisson, S. A. (2016). Modelling extremes using approximate bayesian computation. Extreme Value Modelling and Risk Analysis, (pp. 281–306).

Everitt, R. G. & Rowińska, P. A. (2021). Delayed acceptance abc-smc. Journal of Computational and Graphical Statistics, 30(1), 55–66.

Fearnhead, P. & Prangle, D. (2012). Constructing summary statistics for approximate bayesian computation: semi-automatic approximate bayesian computation. Journal of the Royal Statistical Society: Series B (Statistical Methodology), 74(3), 419–474.

Figueiredo, R., Nunes, P., & Brito, M. C. (2018). Multiyear calibration of simulations of energy systems. Energy, 157, 932–939.

Filippi, S., Barnes, C. P., Cornebise, J., & Stumpf, M. P. (2013). On optimality of kernels for approximate bayesian computation using sequential monte carlo. Statistical applications in genetics and molecular biology, 12(1), 87–107.

Friel, N. & Pettitt, A. N. (2008). Marginal likelihood estimation via power posteriors. Journal of the Royal Statistical Society Series B: Statistical Methodology, 70(3), 589–607.

Fu, Y.-X. & Li, W.-H. (1997). Estimating the age of the common ancestor of a sample of dna sequences. Molecular biology and evolution, 14(2), 195–199.

Gutmann, M. U., Dutta, R., Kaski, S., & Corander, J. (2018). Likelihood-free inference via classification. Statistics and Computing, 28, 411–425.

Hamilton, G., Currat, M., Ray, N., Heckel, G., Beaumont, M., & Excoffier, L. (2005). Bayesian estimation of recent migration rates after a spatial expansion. Genetics, 170(1), 409–417.

Harrison, J. U. & Baker, R. E. (2020). An automatic adaptive method to combine summary statistics in approximate bayesian computation. PloS one, 15(8), e0236954.

Joyce, P. & Marjoram, P. (2008). Approximately sufficient statistics and bayesian computation. Statistical applications in genetics and molecular biology, 7(1).

Khazeiynasab, S. R. & Qi, J. (2021). Generator parameter calibration by adaptive approximate bayesian computation with sequential monte carlo sampler. IEEE Transactions on Smart Grid, 12(5), 4327–4338.

Lenormand, M., Jabot, F., & Deffuant, G. (2013). Adaptive approximate bayesian computation for complex models. Computational Statistics, 28(6), 2777–2796.

Lintusaari, J., Gutmann, M. U., Dutta, R., Kaski, S., & Corander, J. (2017). Fundamentals and recent developments in approximate bayesian computation. Systematic biology, 66(1), e66–e82.

Luciani, F., Sisson, S. A., Jiang, H., Francis, A. R., & Tanaka, M. M. (2009). The epidemiological fitness cost of drug resistance in mycobacterium tuberculosis. Proceedings of the National Academy of Sciences, 106(34), 14711–14715.

Marin, J.-M., Pudlo, P., Robert, C. P., & Ryder, R. J. (2012). Approximate bayesian computational methods. Statistics and Computing, 22(6), 1167–1180.

Marjoram, P., Molitor, J., Plagnol, V., & Tavaré, S. (2003). Markov chain monte carlo without likelihoods. Proceedings of the National Academy of Sciences, 100(26), 15324– 15328.

Nabavi, S. A., Saluz, U., Wolf, M., & Geyer, P. (2023). Hvac systems performance modeling using component-based machine learning. In Building Simulation 2023, volume 18 (pp. 1149–1157).: IBPSA.

Neal, R. M. (1996). Sampling from multimodal distributions using tempered transitions. Statistics and computing, 6, 353–366.

NOAA (2013). National temperature and precipitation analysis. https://www.ncei.noaa.gov/access/monitoring/monthly-report/national/201213.

Peters, G. W., Fan, Y., & Sisson, S. A. (2012). On sequential monte carlo, partial rejection control and approximate bayesian computation. Statistics and Computing, 22(6), 1209– 1222.

Prangle, D. (2017). Adapting the abc distance function. Bayesian Analysis, 12(1), 289–309.

Prangle, D. (2018). Summary statistics. In Handbook of approximate Bayesian computation (pp. 125–152). Chapman and Hall/CRC.

Prangle, D., Everitt, R. G., & Kypraios, T. (2018). A rare event approach to high-dimensional approximate bayesian computation. Statistics and Computing, 28, 819–834.

Pritchard, J. K., Seielstad, M. T., Perez-Lezaun, A., & Feldman, M. W. (1999). Population growth of human y chromosomes: a study of y chromosome microsatellites. Molecular biology and evolution, 16(12), 1791–1798.

Rees, M., Childs, D. Z., Metcalf, J. C., Rose, K. E., Sheppard, A. W., & Grubb, P. J. (2006). Seed dormancy and delayed flowering in monocarpic plants: selective interactions in a stochastic environment. The American Naturalist, 168(2), E53–E71.

Rodrigues, G. S., Francis, A. R., Sisson, S. A., & Tanaka, M. M. (2018). Inferences on the acquisition of multi-drug resistance in mycobacterium tuberculosis using molecular epidemiological data. In Handbook of Approximate Bayesian Computation (pp. 481–511). Chapman and Hall/CRC.

Salguero-Gomez, R., Siewert, W., Casper, B. B., & Tielbörger, K. (2012). A demographic approach to study effects of climate change in desert plants. Philosophical Transactions of the Royal Society B: Biological Sciences, 367(1606), 3100–3114.

Sato, J. (2023). Hand-eye calibration via linear and nonlinear regressions. Automation, 4(2), 151–163.

Schälte, Y., Alamoudi, E., & Hasenauer, J. (2021). Robust adaptive distance functions for approximate bayesian inference on outlier-corrupted data. bioRxiv.

Schälte, Y. & Hasenauer, J. (2022). Informative and adaptive distances and summary statistics in sequential approximate bayesian computation. bioRxiv.

Schulz, E., Speekenbrink, M., & Krause, A. (2018). A tutorial on gaussian process regression: Modelling, exploring, and exploiting functions. Journal of Mathematical Psychology, 85, 1–16.

Scranton, K., Knape, J., & de Valpine, P. (2014). An approximate bayesian computation approach to parameter estimation in a stochastic stage-structured population model. Ecology, 95(5), 1418–1428.

Siciliano, B., Khatib, O., & Kröger, T. (2008). Springer handbook of robotics, volume 200. Springer.

Sisson, S. & Fan, Y. (2018). Abc samplers. Handbook of approximate bayesian computation, (pp. 87–123).

Sisson, S. A., Fan, Y., & Beaumont, M. A. (2018). Overview of abc. In Handbook of approximate Bayesian computation (pp. 3–54). Chapman and Hall/CRC.

Sisson, S. A., Fan, Y., & Tanaka, M. M. (2009). Correction: Sequential monte carlo without likelihoods. Proceedings of the National Academy of Sciences, 106(39), 16889–16890.

Tavaré, S., Balding, D. J., Griffiths, R. C., & Donnelly, P. (1997). Inferring coalescence times from dna sequence data. Genetics, 145(2), 505–518.

Tishkoff, S. A., Varkonyi, R., Cahinhinan, N., Abbes, S., Argyropoulos, G., Destro-Bisol, G., Drousiotou, A., Dangerfield, B., Lefranc, G., Loiselet, J., et al. (2001). Haplotype diversity and linkage disequilibrium at human g6pd: recent origin of alleles that confer malarial resistance. Science, 293(5529), 455–462.

Toni, T., Welch, D., Strelkowa, N., Ipsen, A., & Stumpf, M. P. (2009). Approximate bayesian computation scheme for parameter inference and model selection in dynamical systems. Journal of the Royal Society Interface, 6(31), 187–202.

Wall, J. D. (2000). A comparison of estimators of the population recombination rate. Molecular Biology and Evolution, 17(1), 156–163.

Wegmann, D., Leuenberger, C., & Excoffier, L. (2009). Efficient approximate bayesian computation coupled with markov chain monte carlo without likelihood. Genetics, 182(4), 1207–1218.

Williams, C. K. & Rasmussen, C. E. (2006). Gaussian processes for machine learning, volume 2. MIT press Cambridge, MA.

Zhu, Z., Christodoulou, M. D., & Steinsaltz, D. (2025). Improving Joint Estimation of Vital Rates in IPMs via Gaussian Processes and ABC. bioRxiv.

